# B cell exhaustion associates with poor response to Bacillus Calmette-Guérin immunotherapy in patients with bladder cancer

**DOI:** 10.64898/2026.04.01.715137

**Authors:** Priyanka Yolmo, Kartik Sachdeva, Alanna Brewer, Sindhuja Pattabhi, Gwenaëlle Conseil, Abdulhameed Abdulhamed, Ashley Griffin, Mitchell Jeffs, Haocheng Yu, David Cook, Roger Li, Sonia V. Del Rincon, Madelyn J Abraham, Christophe Goncalves, Lars Dyrskjøt, Trine Strandgaard, Sia Viborg Lindskrog, Amir Horowitz, Peter C. Black, Morgan E. Roberts, David M. Berman, D. Robert Siemens, Madhuri Koti

## Abstract

The majority of patients treated with Bacillus Calmette–Guérin (BCG) immunotherapy, for non-muscle invasive bladder cancer (NMIBC), experience early recurrence due to pre-existing mucosal immune dysfunction. Since B cell are mucosal immune sentinels, we characterized the systemic and local B cell responses in 45 patients with NMIBC. Expansion of circulating atypical B cells (ABCs) following repeated BCG instillation, expanded IgG autoantibody repertoire, progressive IgG reactivity against BCG antigens, and higher tumor IgG deposition, were features of patients who recurred early. Integrated spatial immunophenotyping and single cell spatial transcriptomic analysis of corresponding tumors revealed increased ABCs within tertiary lymphoid structures, and co-localization with PD-1⁺ B cells, regulatory T cells, and CD163⁺ macrophages. Independent validation in two patient cohorts (total n = 409), revealed a significant association between high expression of the ABC specific, *FCRL5*, and poor outcomes. Our study identifies ABCs as key mediators of poor response to BCG in patients with high-risk NMIBC.

## Introduction

Non-muscle invasive bladder cancer (NMIBC) is the most common urological malignancy in females and males, accounting for ∼75% of newly diagnosed bladder cancer cases[1, 2][1, 2]. Standard management for high-grade NMIBC consists of transurethral resection of the bladder tumor (TURBT) followed by repeated intravesical instillation of Bacillus Calmette–Guérin (BCG) immunotherapy over a period of 1-3 years[3, 4]. Despite its established efficacy, durable anti-tumor immune response is not achieved in more than half of the patients receiving adequate doses (5 of induction and 2 of maintenance) of BCG[4–6]. Patients experiencing early recurrence and/or disease progression often require radical cystectomy posing substantial implications on quality of life[7]. Newer second-line treatments for a BCG unresponsive disease, confer a modest response, and also do not provide a durable benefit. Understanding the immunological determinants of immunological responsiveness to BCG immunotherapy is therefore imperative for improved management and incorporation of newer therapeutics in high-risk NMIBC.

BCG exerts its therapeutic effects through a coordinated local and systemic immune activation, encompassing both innate and adaptive immune activation leading to the recruitment of cytotoxic lymphocytes to the bladder mucosa[8–10]. BCG vaccination is also known to induce a long-lived memory B cell response and influence T follicular helper cell differentiation[11–16]. While the role of innate immune reprogramming and T cell–mediated cytotoxicity has been extensively studied[17–21], the role of B cells and humoral immune response to BCG exposure remains poorly defined in patients with NMIBC. This gap is notable given that BCG is administered directly into the bladder lumen, where local immune surveillance involves B cells homing to the mucosa, immunoglobulin (Ig) deposition, and formation of ectopic lymphoid aggregate/tertiary lymphoid structures (TLS) in response to persistent inflammatory stimuli of microbial or non-microbial origin. Additionally, our prior studies demonstrated that increased intra-tumoral B cell density correlated with poor clinical outcomes in high-risk NMIBC[22, 23], underscoring the need to better characterize B cell-specific immune responses in the context of BCG response.

Emerging evidence across chronic inflammatory diseases and solid tumors has identified a phenotypically and transcriptionally distinct population of B cells termed age-associated B cells/double negative type 2/atypical B cells (ABCs)[24–29]. ABCs expand during biological aging, under conditions of persistent antigen stimulation, sustained toll-like receptor (TLR7/9) signaling, and exposure to interferon (IFN)-γ, and interleukin (IL)-21[30]. Their emergence reflects a deviation from canonical B cell maturation, characterized by attenuated B cell receptor (BCR) signaling, expression of the surface markers CD11c and FCRL5, and reduced immune activation capacity[31–33]. Elevated ABC frequencies have been linked to impaired humoral immunity and resistance to immunotherapies across several solid tumors[24, 25, 27, 34]. Using a carcinogen-induced aging murine model of NMIBC, we previously demonstrated that repeated BCG instillations drive systemic and local expansion of ABCs leading to a pre-treatment exhausted immune environment[9], establishing the mechanistic association between ABC expansion, impaired response to BCG, and disease progression.

NMIBC predominantly affects older adults[35], who exhibit reduced naïve lymphocyte pools, decreased clonal diversity, and accumulation of terminally differentiated, memory-like senescent and exhausted immune subsets[36–41]. Recent reports demonstrate that ABCs drive CD4 and CD8 T cells to senescent states via both MHC dependent and independent mechanisms[30]. Our previous report in aging carcinogen induced murine models of NMIBC and patient tumors demonstrated that B cell dominant tumor adjacent (TA)-TLS in pre-treatment tumors from BCG non-responders are enriched for cells expressing immune exhaustion-associated proteins[9, 42]. Collectively, findings to date are suggestive of a potential compounding effect of ABCs on effector T cell senescence and chronic mucosal inflammation in creating a local and systemic immune landscape permissive to ABC accumulation and dysregulated B cell maturation during BCG therapy in patients who experience early recurrence following treatment.

Herein, we performed an integrated analysis of local and systemic humoral immune responses associated with repeated BCG exposure via longitudinal profiling of tumor specimens and corresponding peripheral blood B cells from patients with high-risk NMIBC. Our findings provide the first evidence that early recurrence following adequate BCG therapy is associated with a pre-existing, ABC-driven local and systemic immune state that is further augmented by repeated instillation of BCG. Importantly, our novel findings contribute to the current state of knowledge on the mode of action of intravesical BCG in NMIBC and position local abundance and systemic expansion of ABCs as potential predictive biomarkers of response to BCG as well as newer immunotherapeutic agents in patients with high-risk NMIBC.

## Results

### Exhausted B cell subsets exhibit distinct spatial enrichment within the tumor-associated niches of BCG non-responders

To build upon our previous findings in murine models of NMIBC and patient tumors delineating the spatial distribution of B cell subsets in the tumor immune microenvironment (TIME) [9, 42], we performed multiplex immunofluorescence (mIF) on pre-BCG whole tumor sections (n=31) from 27 patients (pre-BCG tumors from 18 responders and paired pre- and post-BCG recurrence tumors from 9 non-responders; **Fig 1A, Suppl. Table S1/Cohort 1**). Custom-designed ABC and T cell exhaustion (T_ex_) antibody panels (8 markers in each panel; **Suppl. Fig. S1A; Suppl. Table S2**) enabled high-dimensional immune profiling of four spatially defined regions across spatial compartments including tumor epithelium (TE), TA-stroma, mature/immature TA-TLS, and lamina propria (LP) (**Fig. 1C; Suppl. Fig. S1A**).

**Figure 1.**
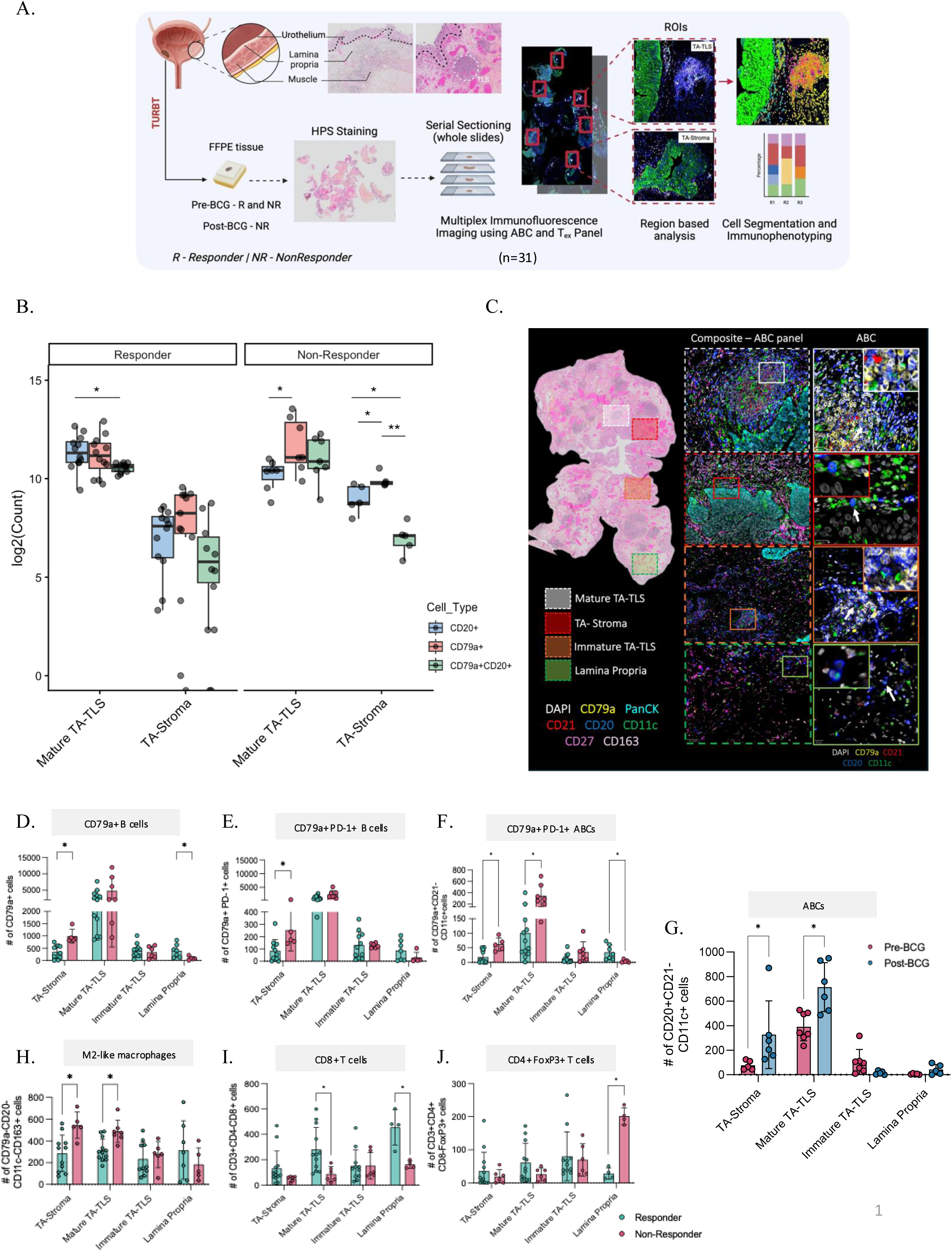
Spatial enrichment of ABCs and CD79a^+^PD-1^+^ B cells within tumor-associated niches define an immunosuppressed microenvironment. Schematic overview of the mIF workflow applied to pre-BCG whole tumor sections responders (n = 18) and paired pre- and post-BCG sections from 9 non-responders. Serial sections underwent HPS staining followed by mIF imaging using the custom ABC and T_ex_ antibody panels. ROIs were defined across four spatially distinct compartments: TA-stroma, mature TA-TLS, immature TA-TLS, and LP, followed by region-based cell segmentation and immunophenotyping (**A**). Box plots showing log₂-transformed counts of CD20^+^, CD79a^+^, and CD79a^+^CD20^+^ double-positive B cells across spatially defined compartments (mature TA-TLS and TA-stroma) in responders and non-responders (**B**). Representative HPS 20x image depicting annotated ROIs (left) across four different compartments (TA-stroma, mature TA-TLS, immature TA-TLS, and LP) along with composite high-magnification insets of the ABC panel (center), and single-channel ABC marker visualization (right), stained for DAPI, CD79a, PanCK, CD21, CD20, CD11c, CD27, and CD163. Arrows indicate representative ABC-phenotype cells (CD79a^+^CD21^−^CD11c^+^; C). Bar plots depicting the total number of CD79a^+^ B cells (**D**), CD79a^+^PD-1^+^ B cells (**E**) and CD79a^+^CD21^−^CD11c^+^ ABCs (**F**) across spatially defined compartments, stratified by treatment response. Bar plots showing the number of CD20^+^CD21^−^CD11c^+^ cells (**G**) across spatial compartments in non-responders pre- and post-BCG. Bar plots depicting the number of CD79a^−^CD20^−^CD11c^−^CD163^+^ M2-like macrophages (**H**). CD3^+^CD4^−^CD8^+^ cytotoxic T cells (**I**), and CD3^+^CD4^+^CD8^−^FoxP3^+^ T_regs_ (**J**) across spatially defined compartments, stratified by treatment response. All bar graphs represent mean ± SD. Statistical comparisons between responders and non-responders across spatial compartments were performed using the Mann–Whitney U test. Longitudinal comparisons within non-responders were performed using the Wilcoxon signed-rank test. Statistical significance thresholds: *\*p < 0.05, **p < 0.01, ***p < 0.001*.

Quantitative spatial analysis revealed both region- and response-associated differences in B cell distribution. In responders, CD79a⁺ and CD20⁺ B cell densities were relatively uniform across all regions (**Fig. 1B, C; Suppl. Fig. S1B**) in contrast to non-responders where preferential accumulation of CD79a⁺ B cells was only observed within the TA-stroma and mature TA-TLS of pre-BCG tumors (**Fig. 1B, C; Suppl. Fig. S1B**). Statistical comparison of annotated regions between responders and non-responders exhibited accumulation of both CD79a^+^ and CD20⁺ B cells in responders within the TA-stroma, while responders demonstrated selective CD20⁺ B cell enrichment within mature TA-TLS (**Fig. 1D**; **Suppl. Fig. S1C**). Furthermore, co-expression analysis revealed that fewer than half of CD79a⁺ and CD20⁺ cells were double positive, underscoring phenotypic heterogeneity among intratumoral B cells. Spatial mapping of PD-1 expression identified significant enrichment of CD79a⁺PD-1⁺ and CD20⁺PD-1⁺ B cells within the TA-stroma of tumors from non-responders (**Fig. 1E; Suppl. Fig. S1D**). In contrast, CD20⁺PD-1⁺ B cells in responders preferentially localized to mature TA-TLS regions (**Suppl. Fig. S1D**). Memory B cell frequencies (CD21⁻/⁺CD27⁺) remained similar across groups and were largely unaffected by BCG treatment in post recurrence tumors (**Suppl. Fig. S1E, F**).

Based on our previous findings in aging mice exposed to bladder specific carcinogen, we next evaluated the spatial distribution of ABCs using a composite classifier (CD79a⁺/CD20⁺CD21⁻CD11c⁺; **Fig. 1C**). ABCs were significantly enriched within the TA-stroma and mature TA-TLS of non-responders in both pre- and post-BCG tumors, whereas no enrichment was observed in immature TA-TLS (**Fig. 1F; Suppl. Fig. S1G**). In responders, ABCs were preferentially localized to the LP in pre-BCG tumors (**Fig. 1F; Suppl. Fig. S1G**). Assessment of tumor sections from BCG non-responders demonstrated a consistent post-treatment increase in ABC density within TA-stroma and mature TA-TLS region (**Fig. 1G; Supplementary Fig. S1J**). Overall, these findings suggest that spatial enrichment of CD79a⁺PD-1⁺ B cells and ABCs within TA stromal niches contributes to the immunosuppressive architecture associated with poor response to BCG.

### Enrichment of cytotoxic T cells and dendritic cells within mature TA-TLS correlates with response to BCG

We next examined whether the spatial profiles of B cells were accompanied by broader remodeling of the myeloid and T cell subsets within the neighboring regions (**Suppl. Fig. S1M**). CD11c^+^ dendritic cells (DCs) were enriched within mature TA-TLS of responders (**Suppl. Fig. S1N**). In contrast, CD163^+^ M2-like macrophages were significantly enriched in the TA-stroma and mature TA-TLS of non-responders (**Fig. 1H**). These spatial distributions were largely unchanged in recurrent tumors from non-responders post adequate BCG therapy (**Suppl. Fig. S2A, B**).

Analysis of T cell subsets (T_ex_ panel; **Suppl. Fig. S2C**) revealed significantly higher total CD3⁺ T cell density within mature TA-TLS regions of pre-BCG tumors from responders compared to non-responders (**Suppl. Fig. 2D**). Cytotoxic T cells (CD3⁺CD4⁻CD8⁺) were selectively enriched in mature TA-TLS and LP regions of responders (**Fig. 1I**), whereas helper T cell (CD3⁺CD4^+^CD8^−^) densities did not differ significantly between groups (**Suppl. Fig. S2E**). PD-1 expression on total T cells and cytotoxic T cells was comparable across response groups (**Suppl. Fig. S2F**); however, PD-1⁺ helper T cells were enriched within mature TA-TLS of responders (**Suppl. Fig. S2G**). Regulatory T cells (T_regs_; CD3⁺CD4⁺CD8⁻FoxP3⁺) were significantly enriched in the LP region of non-responders (**Fig. 1J; Suppl. Fig. S1M**).

Comparison of pre- and post-BCG tumors in non-responders revealed an overall increase in T cell densities across compartments following repeated BCG exposure (**Suppl. Fig. S1M**). Helper T cells preferentially expanded within mature TA-TLS, while cytotoxic T cells irrespective of PD-1 expression accumulated within the TA-stroma (**Suppl. Fig. S2H-K**). However, PD-1⁺ helper T cells and T_reg_ cell densities did not significantly change between timepoints (**Suppl. Fig. S2J, L**), suggestive of repeated BCG driving spatially organized immunosuppressive niches linked to early recurrence in patients with NMIBC.

### Spatial profiling reveals distinct immune neighborhoods and epithelial proximity of ABCs in non-responders

Having established that ABCs are enriched in the bladder TIME of BCG non-responders and their increase in tumors post-BCG recurrence, we next investigated their spatial organization relative to other immune populations. To achieve this, we profiled a subset of cohort 1 (12 whole tumor sections; 3 responders + 5 non-responders (pre-BCG and post-1^st^ recurrence); **Fig. 2A**, **Suppl. Table S1**) using an expanded mIF panel of 25 immune cell functional state markers using the PhenoCycler Fusion platform. This high-plex phenotyping approach enabled simultaneous annotation of diverse immune and functional markers and facilitated comprehensive spatial neighborhood analysis (**Fig. 2A**). Rather than selecting discrete smaller ROIs from the fragmented tissue from piecemeal TURBT tissues, sections with intact TE and associated stroma/LP (average number of cells ∼60,000) were delineated for analysis to minimize selection bias and capture global tissue architecture (**Fig. 2A**). Cell phenotyping was performed using a supervised classifier, with ABCs defined as CD79a^+^CD21^−^CD11c^+^ cells. The CD79a marker provided broader B cell coverage than CD20, supporting its use for ABC detection in this dataset.

**Figure 2.**
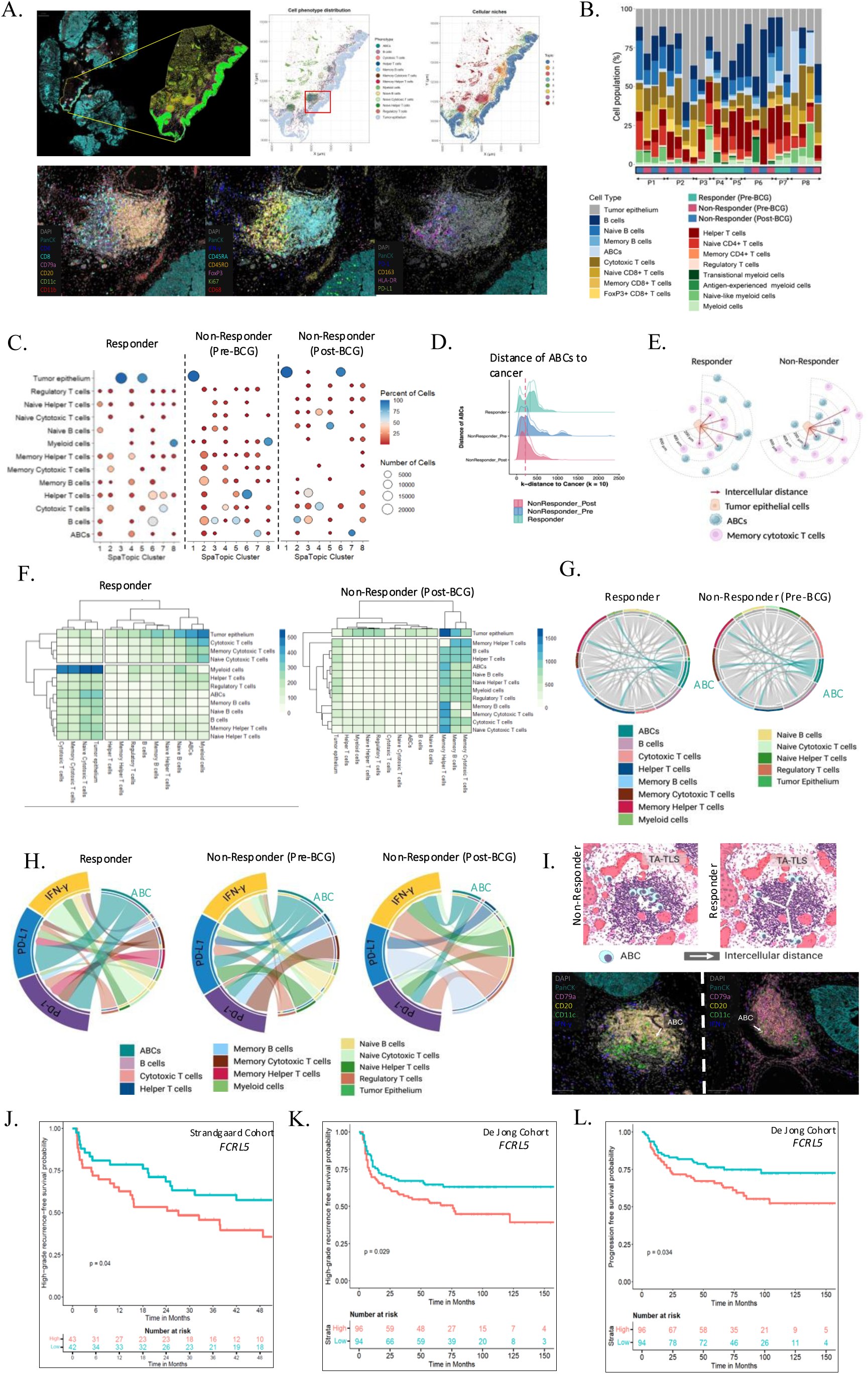
Spatial enrichment of IFN-γ^+^ ABCs within tumor epithelial niches and TA-TLS cores defines an immunosuppressed microenvironment in BCG non-responders, and elevated FCRL5 expression associates with poor clinical outcomes in patients with NMIBC. Representative whole-tumor section image acquired on the PhenoCycler Fusion platform (left), with projected cell phenotype distribution map (center) and spatial niche/cellular neighborhood map (right) illustrating the global immune architecture across the tumor section (**A**). Representative high-plex mIF images of whole tumor sections stained with 25 markers acquired on the PhenoCycler Fusion platform (n = 12; **A**). Stacked bar plots depicting the proportional distribution of annotated immune cell populations across individual patient samples (P1–P8), stratified by treatment response group **(B).** Dot plots depicting spatial domain composition across annotated cell types in responders (n = 3), non-responders (pre-BCG; n = 5 and post-BCG recurrence; n = 4). Dot size reflects the number of cells; color intensity reflects the percentage of cells within each spatial domain **(C).** Density plot illustrating the k-nearest neighbor (k = 10) distance distribution of ABCs to tumor epithelial cells across response groups (responder, non-responder pre-BCG, non-responder post-BCG), demonstrating significantly closer epithelial proximity of ABCs in non-responders **(D).** Schematic figure depicting spatial co-localization relationships between ABCs, tumor epithelial cells, and memory cytotoxic T cells in responders (left) and non-responders (right). Line length reflects intercellular distance **(E).** Heatmap with hierarchical clustering depicting pairwise spatial co-localization scores between annotated immune and epithelial cell populations in responders and non-responders post-BCG **(F).** Color intensity reflects the degree of spatial co-localization; clustering reveals distinct immune neighborhood architecture across response groups. Chord diagrams illustrating predicted cell–cell interaction networks centered on ABCs across annotated immune populations in responders and non-responders pre-BCG **(G**). Arc width reflects interaction frequency or strength between connected cell populations. Chord diagrams depicting the distribution of IFN-γ, PD-L1, and PD-1 expression across annotated immune and epithelial cell populations in responders, non-responders pre-BCG, and non-responders post-BCG (**H**). Arc width reflects the proportion of cells expressing each marker within each annotated population. Representative schematic of HPS-stained (top) and mIF composite images (bottom; DAPI, CD79a, CD20, CD11c) illustrating the distinct spatial localization of ABCs relative to TA-TLS in responders and non-responders. In responders, ABCs localize to the TLS periphery consistent with organized germinal center architecture; in non-responders, ABCs accumulate within TLS cores occupying intrafollicular regions (**I**). Kaplan–Meier survival curves depicting RFS stratified by *FCRL5* gene expression level (high, intermediate, low) in two independent cohorts of patients with high-risk NMIBC (n =126 [*Strandgaard Cohort*] and n = 283 [*de Jong Cohort[44]*]; **J-L**).

Cytotoxic T cells and myeloid cells were relatively enriched in non-responders, whereas helper T cells were more abundant in responders (**Fig. 2B; Suppl. Fig. S3A**). Tumors from non-responders exhibited a reduction in memory B cells, a pattern that persisted following repeated BCG exposure (**Fig. 2C; Suppl. Fig. S10A**). Spatial domain analysis integrating proximity-based neighborhood enrichment and cell-type co-occurrence within shared domains revealed ABC-containing domains were enriched for helper T cells, myeloid cells, and other B cell populations, consistent with integration into an antigen-presenting and T cell supportive immune niche in the tumors from responders (**Fig. 2C**). In contrast, ABC-containing domains in non-responders were characterized by enrichment of cytotoxic T cells and B cells with limited co-enrichment of helper T cells or myeloid populations, indicating a distinct immune architecture associated with poor response to BCG (**Fig. 2C**). Within epithelial-associated spatial domains, an immune cell-devoid niche was observed in non-responders in contrast to tumors from responders where cytotoxic T cell enrichment was observed (**Fig. 2C**).

To quantify epithelial proximity, k-nearest neighbor analysis was performed. ABCs were located near the TE in non-responders compared to that of responders (**Fig. 2D, E; Suppl. Fig. S3B**). In the TE of responders, cytotoxic T cells, helper T cells, and naïve T cell subsets co-localized, while remaining spatially separated from ABCs, myeloid cells, memory B cells, and memory T cells (**Fig. 2E, F**). In contrast, pre-BCG tumors from non-responders exhibited epithelial co-localization of ABCs, T_regs_, naïve B cells, cytotoxic T cells, and memory cytotoxic T cells, and lack of memory B and T cell populations (**Fig. 2F; Suppl. Fig. S3C**). Cell–cell interaction analysis mirrored these spatial patterns. In responders, ABC interactions were predominantly observed with helper T cells and other B cell populations, whereas ABCs in tumors from non-responders displayed increased interactions with cytotoxic T cells in addition to helper T and myeloid subsets (**Fig. 2G; Suppl. Fig. S3D**).

We next examined expression of PD-1, PD-L1 immune checkpoints, and IFN-γ, across the annotated cell phenotypes (**Fig. 2H**). ABCs consistently expressed PD-1 and PD-L1 in tumors from both responders and non-responders, indicative of their exhausted phenotype within the bladder TIME. Expression patterns of IFN-γ displayed a distinct distribution, with no detectable IFN-γ observed in responder tissues, whereas the tumors from non-responders (5/9) showed high IFN-γ expression in the TIME (**Fig. 2H**). Helper T cells in non-responders also exhibited increased PD-L1 expression, suggesting enhanced engagement with antigen-presenting cells within the TIME. PD-L1 expression was also observed on TE from post-BCG non-responders, consistent with induction of immune checkpoint signaling following repeated inflammatory stimulation (**Fig. 2H**).

Qualitative spatial mapping revealed distinct patterns of ABC localization relative to TA-TLS. In responders, ABCs were predominantly located at the periphery of mature TLS, consistent with organized GC architecture (**Fig. 2I**). In contrast, non-responders displayed dense accumulation of ABCs within TLS core regions, occupying intrafollicular regions (**Fig. 2I**). Collectively, these findings demonstrate that ABCs in tumors from BCG non-responders occupy distinct spatial niches, display altered immune interaction networks and exhausted states within an IFN-γ–dominant inflammatory microenvironment and potentially contribute to a tumor permissive immune environment.

Given the known ABC-specific expression of *FCRL5*, we evaluated the association between its expression patterns and survival outcomes using tumor whole transcriptome profiles of two independent cohorts (Strandgaard cohort*[43],* n=126; de Jong et al.,[44] n=283) of BCG treated patients with high-risk NMIBC. While increased expression of *FCRL5* in pre-treatment tumors from both cohorts (total n=409 patients) associated with significantly shorter high-grade recurrence-free survival (RFS), a significant association with shorter progression free survival (PFS) was observed only in the deJong cohort (**Fig. 2J-L**). These findings establish elevated tumor *FCRL5* expression as a potential predictive biomarker associated with poor prognosis in patients undergoing BCG therapy.

### Single-cell spatial transcriptomic profiling of the bladder TIME reveals enrichment of B cell and plasma cell-associated signatures in BCG non-responders

To further resolve the cellular states underlying the spatial immune architecture identified by mIF, we performed high-resolution single cell spatial transcriptomic profiling of 5,000 genes using the Xenium In Situ platform in a subset of whole tumor sections. Areas of interest selection from a subset of six whole tumor sections from two responders (pre-BCG) and two non-responders (pre-BCG and post-BCG recurrence) was guided by 25-plex mIF based immune cell spatial profiles; **Suppl. Table S1/Cohort 1 and 2**). This approach enabled single-cell-resolved transcriptomic characterization precisely within histologically annotated regions previously defined by mIF, allowing integration of spatial protein-level architecture with *in situ* transcriptional profiles (**Fig. 3A**). Unsupervised clustering of the Xenium based transcriptional profiles identified 12 transcriptionally distinct cell clusters representing epithelial, endothelial, immune, and stromal populations (**Fig. 3B, C; Suppl. Fig. S4A**). Transcripts associated with plasma cells, B cells, myeloid cells, and granulocytes were higher in the tumors from non-responders relative to responders (**Fig. 3C; Suppl. Fig. S4B**).

**Figure 3.**
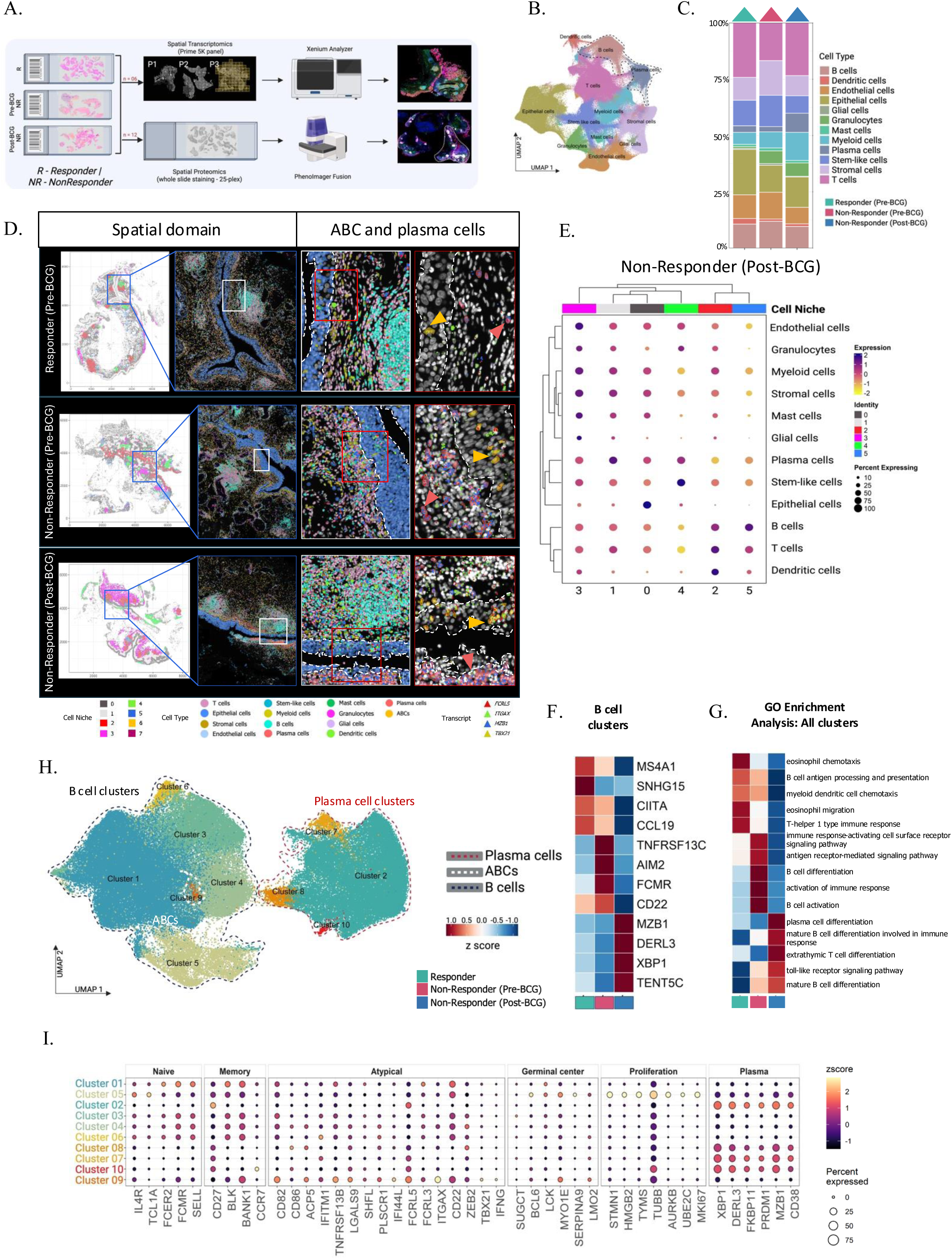
Spatial transcriptomic profiling reveals enrichment of ABC and plasma cell transcripts within the bladder TIME of BCG non-responders. Schematic overview of the spatial multi-omic workflow applied to 6 FFPE bladder whole tumor sections from responders (pre-BCG; n = 2) and non-responders (pre-BCG; n = 2, post-BCG; n = 2). Spatial transcriptomics was performed using the Xenium In Situ platform targeting ∼5,000 genes (Prime 5K panel), and spatial proteomics was performed using whole-slide staining with a 25-plex antibody panel imaged on the PhenoImager Fusion platform on whole tumor sections from 12 patients (**A**). UMAP visualization of unsupervised clustering of the integrated Xenium dataset identifying 12 transcriptionally distinct cell populations, encompassing epithelial, endothelial, immune, and stromal lineages (**B**). Stacked bar plots depicting the proportional distribution of annotated cell types across treatment response groups (responder pre-BCG, non-responder pre-BCG, and non-responder post-BCG) (**C**). Representative spatial domain maps (left) and high-magnification composite images highlighting ABC and plasma cell transcriptional signals (right) across tumor sections from responders (pre-BCG), non-responders (pre-BCG), and non-responders (post-BCG). Annotated transcripts include *FCRL5* (ABCs), *ITGAX* (ABCs), *MZB1* (plasma cells), and *TBX21* (ABC-associated transcription factor). Arrowheads indicate representative cells expressing ABC-and plasma cell-associated transcripts within spatially defined niches (**D**). Dot plot depicting scaled average expression and percentage of cells expressing cell-type marker genes across identified spatial niches in non-responder (post-BCG) tumors. Dot size reflects the percentage of expressing cells; color intensity reflects scaled average expression (**E**). Heatmap displaying z-scored expression of selected B cell and plasma cell-associated marker genes across treatment response groups (responder, non-responder pre-BCG, non-responder post-BCG; **F**). Heatmap showing GO biological process enrichment scores across all identified cell clusters, stratified by treatment response group. Color intensity reflects normalized enrichment score (**G**). UMAP visualization of sub-clustered B cell and plasma cell populations identifying 10 transcriptionally distinct clusters, annotated as naïve/memory-like B cells, plasma cells, GC B cells, and ABCs (**H**). Dot plot displaying scaled average expression and percentage of cells expressing canonical marker genes across B cell subclusters, spanning naïve, memory, atypical, GC, proliferating, and plasma cell transcriptional programs. Dot size reflects the percentage of expressing cells; color intensity reflects scaled z-scored expression (**I**).

To further determine how these cell types were spatially organized within the TIME, we performed niche-based analysis to identify cellular communities that co-localize within the tissue (**Fig. 3D, E; Suppl. Fig. S4C, D**). Each tumor section displayed multiple spatial niches with variable cellular composition, reflecting inter-patient heterogeneity. Across samples, T cells consistently co-localized with DCs within shared niches. Dominant expression of B cell-associated genes was detected across two to three distinct niches, suggesting their presence within TLS-like regions, LP compartments, and TA-stromal areas (**Fig. 3D, E; Suppl. Fig. S4C, D**), consistent with patterns observed by mIF. Myeloid cell abundance positively correlated with stromal cell representation within these spatial niches.

Differential expression analysis of the B cell and plasma cell cluster across different groups revealed distinct transcriptional programs associated with treatment response (**Fig. 3F**). Post-BCG recurrence tumors from non-responders exhibited elevated expression of plasma cell-associated transcripts including *XBP1* and *MZB1*, consistent with plasma cell differentiation. Increased expression of *CD22, AIM2,* and *TNFRSF13C (BAFF-R)* was observed in the pre-BCG tumors of non-responders. In contrast, tumors from the two responders showed increased expression of *CCL19* and *CIITA*, genes associated with antigen presentation and regulation of MHC class II expression, indicative of enhanced adaptive immune priming.

GO enrichment analysis of all the clusters across different treatment response groups further revealed enrichment of pathways associated with B cell activation and differentiation as well as TLR signaling in tumors from non-responders, whereas those from responders were enriched for pathways related to DC chemotaxis and Th1-type immune responses (**Fig. 3G**). Consistently, Reactome pathway analysis revealed increased cytokine and IFN signaling in tumors from non-responders (**Suppl. Fig. S4E**).

To further define B cell heterogeneity within the bladder TIME, we performed sub-clustering of the combined B cell subsets and transcriptionally distinct plasma cell populations. This analysis revealed ten transcriptionally distinct clusters (**Fig. 3H; Suppl. Fig. S4F**). Based on the canonical markers, clusters 1,3, 4, and 6 represented a mix of naïve and/or memory-like B cells. Cluster 2, 7, 8, and 10 corresponded to plasma cells expressing *MZB1* and *XBP1,* while cluster 5 exhibited features of GC B cells with expression of *BCL6* and *LCK.* Notably, cluster 9 displayed a transcriptional signature consistent with ABCs, characterized by expression of *ITGAX,* and *FCRL5* (**Fig. 3H, I**). Comparative analysis between groups revealed enrichment of ABC-associated signatures in tumors from non-responders (pre- and post-BCG), whereas tumors from BCG responders contained relatively few ABC and plasma-cell associated transcripts (**Suppl. Fig. S4G**).

### Spatial organization of ABCs within epithelial niches associates with distinct epithelial transcriptional states in tumors from BCG non-responders

To determine the spatial distribution of B cell subsets, the annotation of the ten B cell subsets was projected back onto the global spatial map of the tumor sections in Xenium explorer. Although spatial proteomic analysis detected ABCs across multiple tumor-associated compartments, including TA-TLS and TA-stromal regions (**Fig. 4A**), ABC-associated transcripts primarily localized within epithelial-adjacent regions (**Fig. 4B**). In particular, cells expressing ABC-associated genes were preferentially localized adjacent to epithelial niches in tumors from non-responders, with strong enrichment of *FCRL5* (**Fig. 4B**). Further sub-clustering revealed ten transcriptionally distinct epithelial clusters based on canonical markers (**Fig. 4C, D, and F**). Epithelial clusters were annotated based on canonical markers representing basal (cluster 3, 4, 5, 8), luminal (cluster 1, 2, 6, 7,9, 10), inflamed/immune-activated (cluster 4, 5), mucus-secreting (cluster 6), proliferative (cluster 10), and angiogenic (clusters 8, 9) epithelial states (**Fig. 4F**).

**Figure 4.**
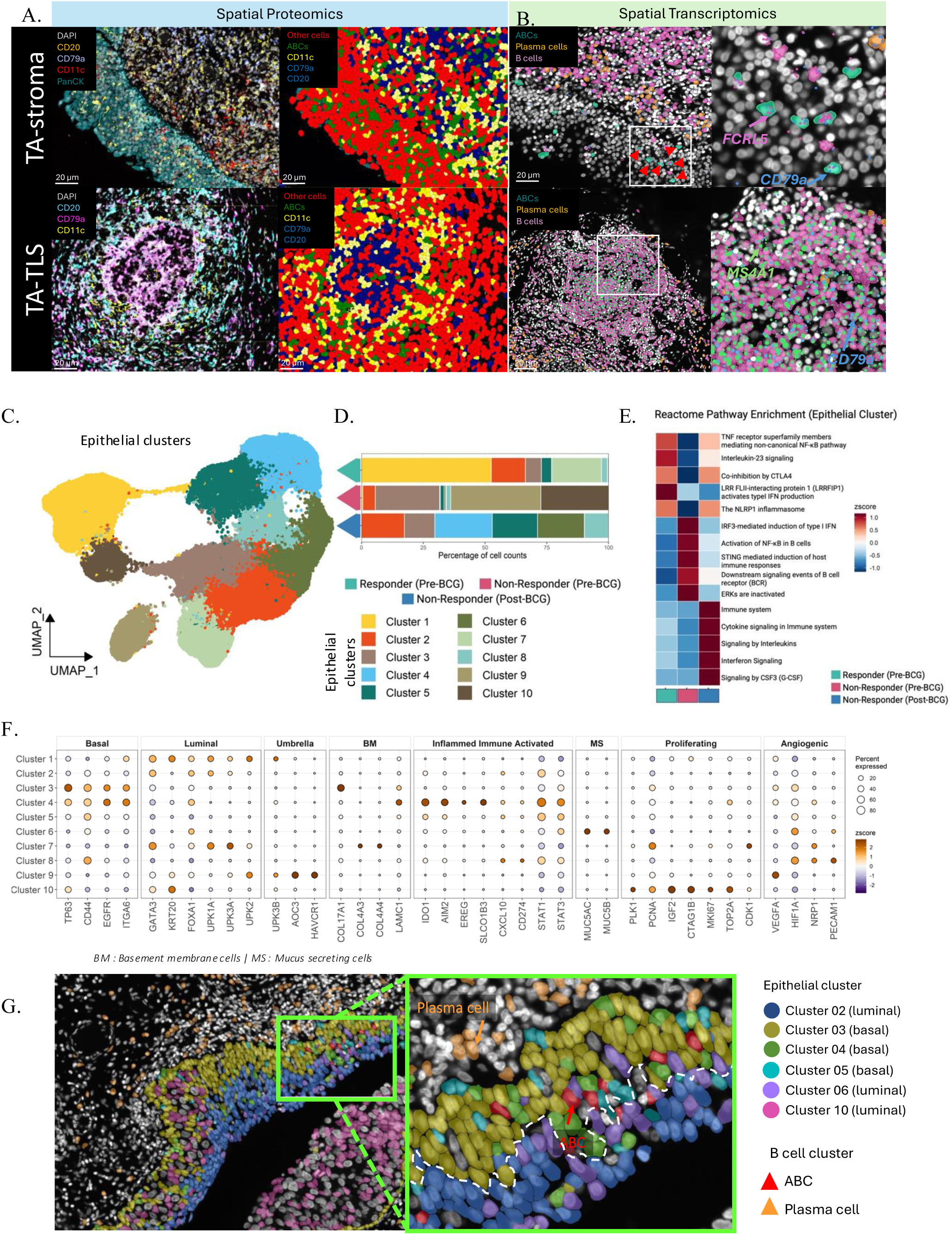
Spatial organization of ABCs within inflammatory basal epithelial niches and elevated tumor FCRL5 expression associate with poor clinical outcomes in patients with NMIBC. Representative mIF images (spatial proteomics) of TA-stroma (top) and TA-TLS (bottom) regions showing composite channel visualization (left; DAPI, CD20, CD79a, CD11c, PanCK) and cell-segmented phenotype maps (right; annotated as ABCs, CD11c⁺ cells, CD79a⁺ cells, CD20⁺ cells, and other cells) (**A**). Representative spatial transcriptomic images from Xenium Explorer 4 displaying the projected distribution of annotated B cell subsets: ABCs, plasma cells, and B cells across tumor sections (left), with high-magnification insets highlighting single-transcript resolution of ABC-associated genes (*FCRL5, CD79A*) and B cell markers (*MS4A1*) within tumor epithelial-adjacent regions (**B**). UMAP visualization of sub-clustered epithelial cells identifying 10 transcriptionally distinct epithelial states (**C**). Stacked bar plots depicting the proportional distribution of epithelial subclusters across treatment response groups (responder pre-BCG (n = 3), non-responder pre-BCG (n = 5), non-responder post-BCG (n = 4) (**D**)). Heatmap showing Reactome pathway enrichment scores across epithelial clusters, stratified by treatment response group (**E**). Dot plot displaying scaled average expression and percentage of cells expressing canonical marker genes used to annotate epithelial subclusters. Dot size reflects the percentage of expressing cells; color intensity reflects scaled z-scored expression (**F**). Representative spatial map from Xenium In Situ profiling illustrating the co-localization of ABC (marked with red arrowheads) and plasma cell (marked with orange arrowheads) transcriptional signatures within basal epithelial clusters across a representative non-responder tumor section (**G**).

Pathway enrichment analysis of epithelial clusters revealed that post recurrence tumors from non-responders were enriched for pathways related to TNF and IFN signaling. Pre-BCG tumors from non-responders showed enrichment of B cell signaling pathways, whereas those from responders exhibited enrichment of TNF receptor and IL-23 signaling (**Fig. 4E**).

Tumors from responders were enriched for luminal epithelial states, whereas those from non-responders exhibited predominance of basal signatures (**Fig. 4D; Suppl. Fig. S4H**). The inflamed epithelial cluster, mucus-producing epithelial cluster and angiogenic basal epithelial states were predominantly detected in post-BCG recurrent tumors. ABCs expressing *FCRL5* were most frequently localized in basal layer (cluster 4, 5; **Fig. 4G**), which showed elevated expression of inflammatory/immune activated genes (*STAT1, STAT3, AIM2, IDO1*; **Fig. 4F**).

### Increase in circulating ABC frequency post-BCG treatment is associated with early recurrence

To determine whether systemic and local expansion of ABCs (CD19^⁺^CD21^⁻/low^CD11c) occur in BCG non-responders, we characterized the corresponding temporal frequencies of circulating ABCs in peripheral blood collected at pre- (n =31), post-1^st^ (n = 25) and post-4^th^ BCG (n = 26) instillation during the induction phase (**Fig. 5A; Suppl. Table S1/Cohort 1 and 2**). Based on the surface expression of CD27, IgD, IgM, CD21, and FCRL5, B cells and ABCs were further resolved into phenotypically distinct subsets (**Fig. 5B; Suppl Fig. S5A, B**).

**Figure 5.**
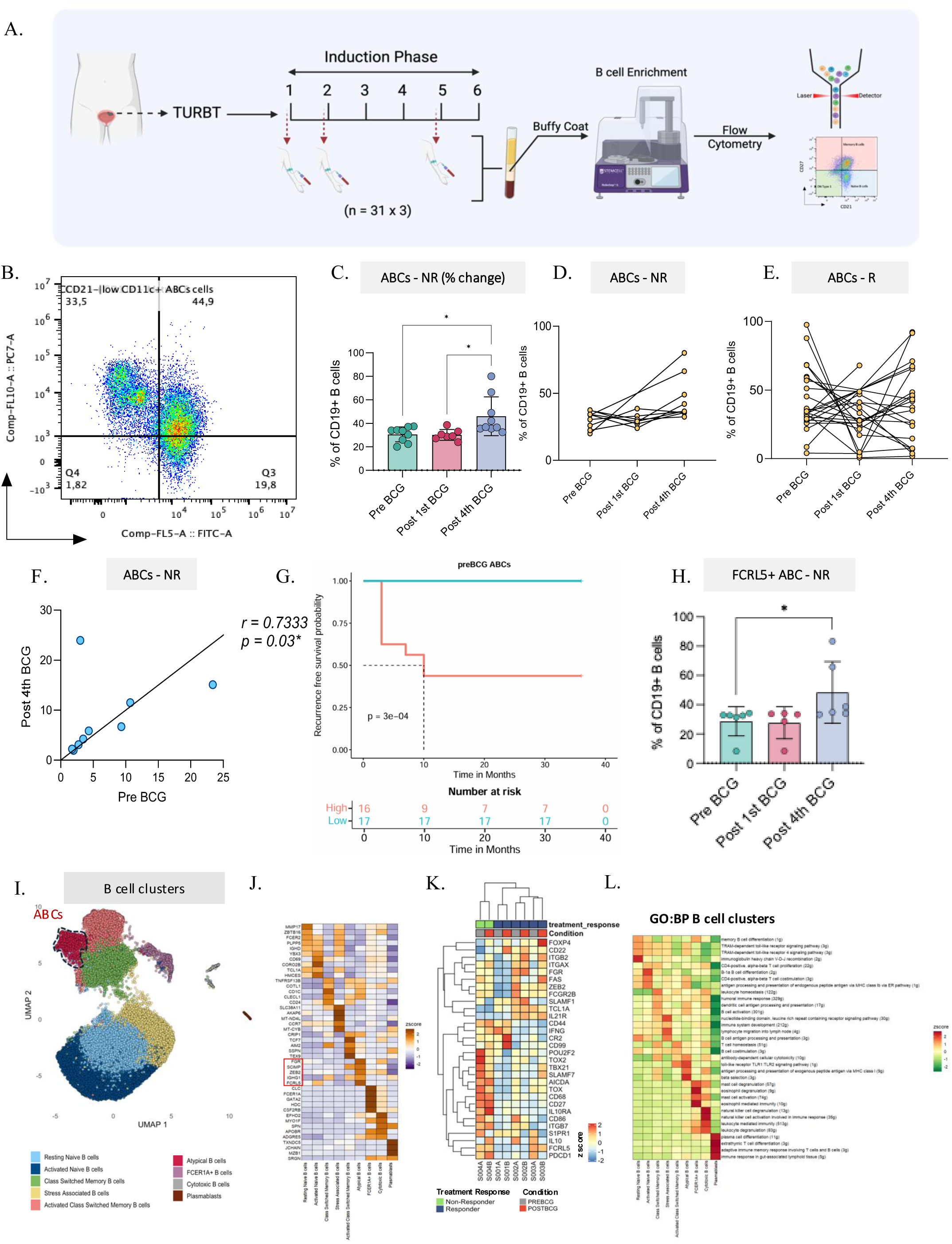
Longitudinal immune profiling reveals expansion of circulating ABCs and differing pre-existing B cell-associated transcriptional enrichment and dominant intercellular communication signatures in BCG non-responders. Schematic overview illustrating the longitudinal profiling of peripheral blood in patients with high-risk NMIBC undergoing BCG induction therapy following TURBT. B cells were enriched from the buffy coat at different timepoints and analyzed by multiparametric flow cytometry (**A**). Gating strategy defining circulating ABCs (CD19⁺ CD21⁻CD11c⁺) (**B**). Bar plot depicting the percentage change in circulating ABCs across treatment timepoints (pre-BCG, post-1^st^ BCG, post-4^th^ BCG) in non-responders (**C**). Paired line plot illustrating longitudinal trajectories of circulating ABCs in non-responders (**D**) and responders (**E**). Spearman correlation plot demonstrating a positive association of ABCs between pre-BCG and post-4^th^ BCG timepoints in non-responders (*r = 0.7333; *p = 0.03;* F). Kaplan-Meier curve depicting RFS stratified by pre-BCG circulating ABC score (high vs. low ABC, defined by median stratification based on multiparametric flow cytometry; **G**). Statistical significance (*p <0.05*) was assessed using the log-rank test. Bar plot showing longitudinal changes in, FCRL5⁺ ABCs (H). UMAP visualization of unsupervised clustering identifying nine transcriptionally distinct B cell populations, with ABCs highlighted in black dotted lines (**I**). Heat map showing z-scored expression of canonical marker genes used for cluster annotation across all identified B cell subsets. (**J**). Heatmap displaying z-scored expression of ABC-associated and exhaustion-related gene signatures across individual patient samples, stratified by treatment response (responder vs. non-responder) and timepoint (pre-BCG vs. post-4^th^ BCG). Hierarchical clustering was applied to both genes and samples (**K**). Heatmap showing pathway-level gene ontology (GO) enrichment scores across annotated B cell subsets, highlighting functional distinctions between naïve, memory, ABC, and effector populations (**L**). All cell frequencies are expressed as a percentage of CD19⁺ B cells. Bar graphs represent mean ± SD. Statistical comparisons across longitudinal timepoints were performed using ordinary one-way ANOVA with Tukey’s post hoc test (**C)**. Significance thresholds: *\*p < 0.05, **p < 0.01, ***p < 0.001, ****p < 0.0001*. Longitudinal timepoints: pre-BCG, post-1^st^ BCG, and post-4^th^ BCG. R, responder (n = 22 x 3); NR, non-responder (n = 9 x 3).

BCG non-responders exhibited a significantly increased frequency of ABCs following the 4^th^ BCG instillation compared to both pre-BCG and post-1^st^ BCG levels (**Fig**. **5C, D****; Suppl Fig. S5A**). In contrast, BCG-responders displayed variable ABC frequencies without significant temporal correlation (**Fig.5E; Suppl. Fig. S6C, D**). Spearman correlation analysis depicted significant positive correlation between circulating ABC profiles at pre- and post-4^th^ BCG timepoints within individual non-responders (*r = 0.7333, p = 0.03*; **Fig. 5F**). In accordance with these results, patients (cohorts 1 and 2) with high pre-BCG systemic ABC frequency, defined using median stratification, exhibited significantly shorter recurrence free survival compared to patients with low ABC frequency (**Fig 5G**).

Consistent with these findings, CD19⁺CD21⁻CD27⁻ double-negative type 1 (DN1) B cells followed a similar expansion pattern in non-responders (**Suppl. Fig. S6E**), but not in responders (**Suppl. Fig. S6F**). Subclassification of ABCs by IgD and CD27 expression further revealed a selective expansion of CD27⁻IgD⁻ DN2 ABCs in non-responders following the 4^th^ BCG instillation, an expansion that was not observed after the 1^st^ instillation (**Suppl. Fig. S6G**). All other ABC subsets remained largely unchanged across both responders and non-responders at both timepoints (**Suppl. Fig. S6H-L**). Although total FCRL5 expression was comparable across timepoints, frequency of post-4^th^ BCG circulating FCRL5⁺ ABCs were significantly higher in non-responders relative to responders (**Fig. 5H; Suppl. Fig. S6M**). These findings are suggestive of a potential influence of repeated exposure to BCG in the systemic expansion of ABCs.

### Differentiation of systemic B cells is distinct between BCG responders and non-responders

Given the divergent ABC trajectories, we next examined whether BCG response was associated with distinct patterns of memory B cell frequencies. Unsupervised FlowSOM clustering revealed a trend towards reduced circulating memory B cells (Pop 2, CD27^+^IgD^+^) post 4^th^ BCG in responders, whereas non-responders retained these populations (**Fig. S6N**). Analysis of IgD and IgM expression further revealed a decline in class-switched B cells (CD19⁺IgD⁻IgM⁻) in responders post-BCG, contrasted by an expansion of this subset in non-responders (**Suppl. Fig. S6O, P**). Stratification into class-switched (CD27^+^IgM^−^IgD^−/+^) and unswitched (CD27^+^IgM^+^IgD^+^) memory B cell compartments revealed that responders exhibited reduced frequencies of class-switched memory B cells accompanied by a reciprocal increase in naïve B cells following BCG induction (**Suppl. Fig. S6Q, R**). In contrast, non-responders showed stable class-switched memory frequencies between pre- and post-4^th^ BCG timepoints (**Suppl. Fig. S6T, M**), alongside a significant reduction in unswitched memory B cells (**Suppl. Fig. S6T, U**). These differential class switching patterns between the two groups suggest that BCG efficacy may require optimal regulation of B cell differentiation.

### Circulating B cells exhibit distinct pre-treatment transcriptional states associated with BCG response

The divergent B cell trajectories observed by flow cytometry, characterized by expansion of ABC and FCRL5⁺ memory populations in non-responders, raised the question of whether the response-specific cellular patterns were pre-configured at the transcriptional level prior to therapy. To address this, we performed single-cell RNA sequencing (scRNA-seq) on enriched peripheral B cells from a subset of four patients before and after the 4^th^ BCG instillation (total 8 enriched B cell samples**; Suppl. Table S1/Cohort 2; Suppl Fig. S7A-C**). Unsupervised clustering of the integrated dataset identified nine transcriptionally distinct B cell populations (**Fig. 5I**).

Cell type annotation using canonical marker genes delineated major B cell populations, including naïve B cells, class-switched memory B cells, ABCs, and plasmablasts (**Fig. 5J**, canonical marker genes listed in **Suppl. Table S3**)[24, 27, 28]. Additional transcriptionally distinct B cell clusters included cytotoxic B cells, *FCER1A⁺* B cells, and stress-associated B cells (**Suppl. Fig. S7D, S7E**). The relative frequencies of these subsets varied substantially between the B cells from 4 patients (**Suppl. Fig. S7E**). ABCs clustered near memory B cell populations while maintaining a transcriptionally distinct identity (**Fig. 5I, J)**.

We next assessed sample-level expression of genes associated with the ABC transcriptional signature and exhaustion-like states. B cells from the patient categorized as the only BCG non-responder exhibited a distinct B cell transcriptional profile, characterized by elevated levels of ABC-associated and chronic activation/exhaustion-related genes[45] (**Fig. 5K**), consistent with the proteomic level findings from our complementary multi-parametric flow cytometry analysis.

To characterize intercellular communication among B cell subsets, we applied CellChat analysis to the single-cell transcriptomic data (**Suppl. Table S3**). CellChat inference indicated enriched autocrine and paracrine signaling involving ABCs (**Supp. Fig. 7F**). Signaling interactions were dominated by pathways associated with immune regulation, antigen responsiveness, and cell-cell adhesion, reflecting the potential role of systemic B cells in shaping the TIME (**Suppl. Fig. S8A-C**).

Within the signaling landscape, ABCs emerged as a particularly communication-active subset (**Suppl. Fig. S8B, C**). ABC-associated interactions were enriched for ligand–receptor pairs involving *CD22*, *PTPRC, FCER2A (CD23), ITGAX,* and *SELL,* molecules implicated in regulation of BCR signaling thresholds, adhesion, and immune modulation under conditions of chronic stimulation (**Suppl. Fig. S7F, S8B, C**). These interactions reflect predicted signaling potential based on ligand–receptor co-expression and do not inform the direct measurements of functional signaling activity.

While class-switched memory B cells preferentially engaged alternative interaction pairs such as *FCER2A–CR2*, ABCs showed stronger associations with integrin- and adhesion-related signaling pathways (**Suppl. Fig. S8D, E**). Communication flow analysis revealed an asymmetric signaling architecture in ABCs, with outgoing signaling patterns resembling naïve B cells and incoming signaling patterns more closely aligned with memory B cells (**Suppl. Fig. S8F-I**).

Pathway enrichment analysis further highlighted functional differences among B cell subsets. Naïve B cells showed enrichment of pathways related to B cell differentiation and antigen recognition, while memory B cell clusters exhibited activation of humoral immune response, B cell co-stimulation, and antibody-dependent cellular cytotoxicity (ADCC; **Fig. 5L**). ABCs shared partial overlap with activated class-switched memory B cells and pathways related to TLR signaling and ADCC (**Fig. 5L**). ABCs also displayed increased enrichment of genes involved in antigen processing and presentation pathways, including those associated with MHC class I presentation (**Fig. 5L**), consistent with an activated yet potentially dysregulated immune phenotype. Together, these findings indicate that pre-existing enrichment of ABC-like and exhausted B cell states potentially prevents effective B cell maturation despite BCG stimulation.

### IgG-skewed antibody profiles and autoantibody diversification characterize poor response to BCG

We next examined whether the changes observed in circulating memory B cell subsets were accompanied by biased Ig class-switching (**Fig. 6A**). Analysis of Ig constant region gene expression across peripheral B cell subsets revealed distinct isotype usage between responders and the non-responder (**Fig. 6B**). Responders showed relatively higher *IGHG3* expression, post-4^th^ BCG instillation, together with variably increased *IGHA1* expression. In contrast, B cells from the non-responder displayed a dominant *IGHG1* and *IGHA2*-biased profile that was evident prior to therapy and sustained following 4 doses of BCG (**Fig. 6B**). Expression of unswitched Ig genes (*IGHM*, *IGHD*) was comparatively limited in the non-responder (**Fig. 6B**), concordant with the reduced frequency of unswitched memory B cells observed by flow cytometry. Quantification of total plasma Ig isotypes using the LegendPlex™ Human Immunoglobulin Isotyping Panel in paired pre- and post-4^th^ BCG samples (n = 20 patients; **Suppl. Table S1/Cohort 1**) revealed no statistically significant differences in total Ig concentrations between responders and non-responders across isotypes (**Suppl. Fig. S9A–E**). Plasma from the non-responders displayed a steady temporal trend toward elevated secreted IgG1 (**Suppl. Fig. S9A**). This trend mirrored the *IGHG1*-biased transcriptional profile and suggests an IgG1-skewed antibody profile in non-responders.

**Figure 6.**
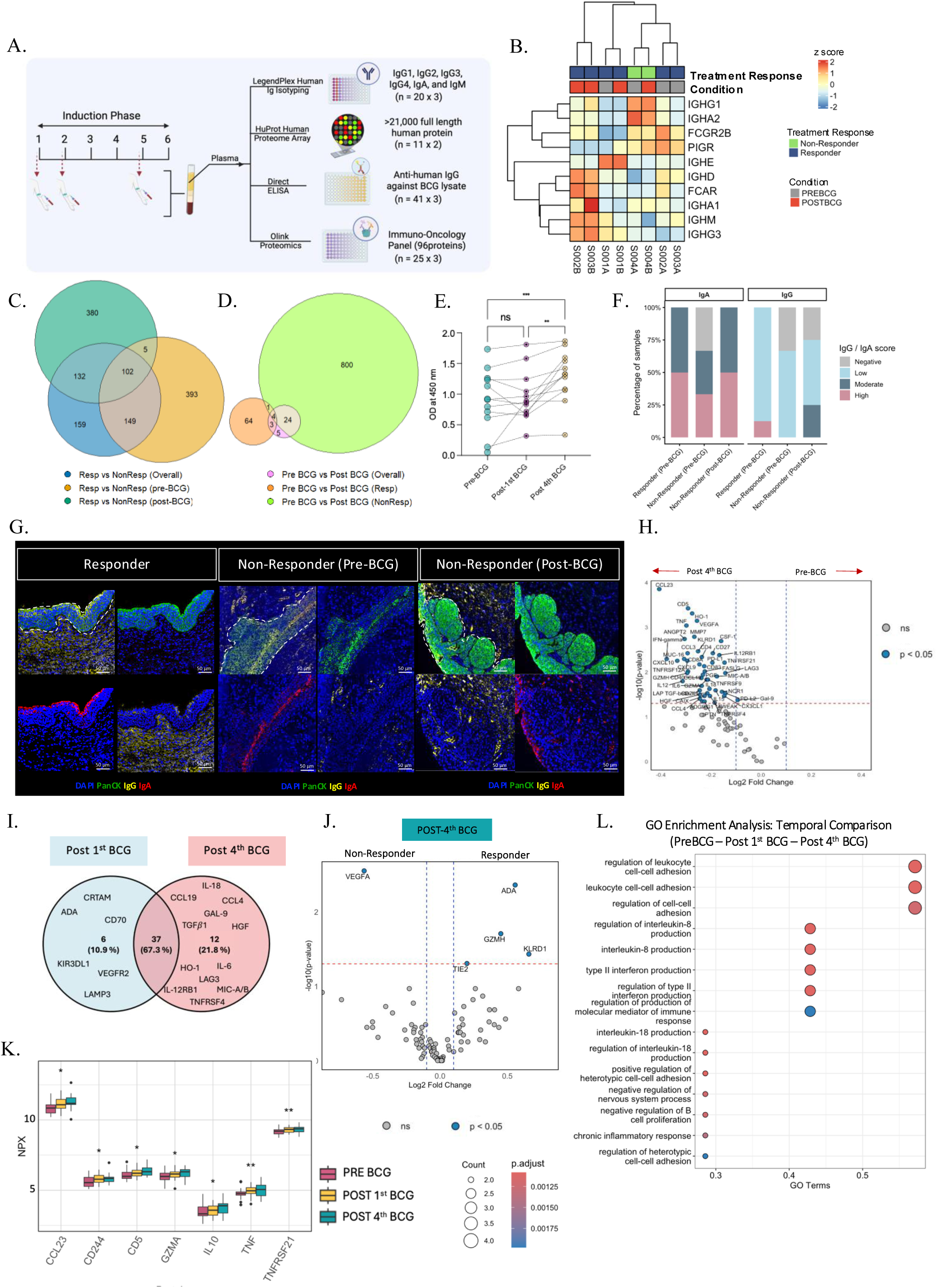
Pre-treatment IgG-skewed humoral immune state and BCG-induced systemic immunosuppression are pronounced in BCG non-responders. Schematic overview of the multi-platform plasma profiling workflow applied to longitudinally collected plasma samples, encompassing LegendPlex™ Human Immunoglobulin Isotyping, HuProt™ Human Proteome Microarray (>21,000 proteins), direct ELISA against BCG lysate, and Olink® Immuno-Oncology proximity extension assay (96 proteins; **A**). Heatmap displaying z-scored expression of Ig constant region genes across individual patient samples (n = 4, pre-BCG and post 4^th^ BCG), stratified by treatment response and timepoint (pre-BCG vs. post-4^th^ BCG). Hierarchical clustering was applied to both genes and samples (**B**). Venn diagrams depicting the overlap of differentially abundant autoantigen-reactive IgG targets between responders (n = 6 x 2) and non-responders (n = 5 x 2) at pre-BCG and post-4^th^ BCG timepoints (**C**) and across treatment timepoints in responders and non-responders separately (**D**). Paired line plot illustrating longitudinal IgG reactivity against BCG lysate (OD at 450 nm) across timepoints (n=12) in non-responders, measured by direct ELISA (**E**). Stacked bar plots depicting semi-quantitative IgG and IgA intensity scores (negative, low, moderate, high) within the TE across responders (n = 8) and non-responders at pre- and post-BCG timepoints (n = 7; **F**). Representative mIF images of bladder tumor sections from responders and non-responders (pre- and post-BCG) stained for DAPI (nuclei), PanCK (epithelial marker), IgG, and IgA (**G**). Volcano plot depicting DEPs identified by Olink® proteomics platform between pre-BCG (n = 27) and post-4^th^ BCG (n = 25) timepoints. Log₂ fold change is shown on the x-axis; −log₁₀(p-value) on the y-axis. Significantly differentially expressed proteins (p < 0.05) are highlighted in blue (**H**). Venn diagram illustrating the overlap of DEPs between post-1^st^ BCG (n = 26) and post-4^th^ BCG (n = 25) timepoints relative to pre-BCG baseline (**I**). Volcano plot comparing plasma protein levels between non-responders (n = 9) and responders (n = 16) at post-4^th^ BCG, with significantly DEPs annotated in blue (**J**). Dot plot showing GO biological process enrichment analysis of DEPs identified in the temporal comparison (pre-BCG → post-1^st^ BCG → post-4^th^ BCG) in non-responders (n = 9 x 3). Dot size reflects gene count; color reflects adjusted p-value (**K**). Box plots depicting normalized protein expression (NPX) levels of significant plasma proteins across longitudinal timepoints (pre-BCG, post-1^st^ BCG, post-4^th^ BCG; n = 25 x 3; **L**). Non-parametric Wilcoxon signed-rank test was performed for paired samples. Friedman test with Dunn’s post hoc correction was used for multiple comparisons. Statistical significance thresholds: *\*p < 0.05, **p < 0.01, ***p < 0.001, ns-not significant*.

Given the chronic nature of the disease, and the potential reactivity against urothelial antigens, we next profiled the qualitative breadth of IgG reactivity using a high-density HuProt^TM^ protein microarray comprising >21,000 conformationally intact human proteins. Analysis of paired plasma samples (n = 11 patients; **Fig. 6A, Suppl. Table S1/Cohort 1**) revealed substantial inter-individual variability. Principal component analysis demonstrated considerable overlap between responders and non-responders across treatment timepoints (**Suppl. Fig. S9F**). Differential abundance analysis identified distinct proteomic profiles between responders and non-responders, with 649 proteins at baseline (pre-BCG) and 619 at the post-fourth timepoint (*log₂ fold change > 1, p < 0.05;* **Fig. 6C, D; Suppl. Fig. S9G**). Longitudinal comparison revealed 829 differentially abundant autoantigens between pre- and post-BCG timepoints of non-responders, whereas responders displayed only 72 autoantigens, with minimal overlap between the groups (**Fig. 6D; Suppl. Fig. S9G; Suppl. Table S4**). Pathway enrichment analysis of the differentially abundant autoantigens in non-responders highlighted biological processes related to antigen processing and presentation, and bacterial invasion of epithelial cells (**Suppl. Fig. S9H**). In contrast, responders showed enrichment of proteins associated with more general cellular and metabolic processes (**Suppl. Fig. S9I**). In summary, these data indicate an expanded pre-BCG IgG autoantibody repertoire in non-responders prior to and following BCG therapy.

Given the expanded autoantigen landscape observed in non-responders, we profiled the altered IgG reactivity against BCG antigens using ELISA. Plasma IgG binding to BCG antigens was measured in plasma samples across the three treatment timepoints (n=41 patients, **Suppl. Table S1/Cohort 3**). Titration experiments identified 1:100 and 1:1,000 plasma dilutions as optimal for detecting non-saturated IgG reactivity (**Suppl. Fig. S9J, K**). Responders exhibited heterogeneous reactivity to BCG antigens without a consistent temporal trend (**Suppl. Fig. S9L, M**), whereas non-responders exhibited a progressive increase in IgG reactivity following repeated BCG exposure (**Fig. 6E; Suppl. Fig. S9N**).

To determine whether systemic Ig profiles were reflected within the corresponding TIME, we assessed IgG and IgA deposition using mIF on tumors from 8 responders (pre-BCG), and 4 non-responders (pre-BCG and post-BCG recurrence; **Suppl. Table S1/Cohort 1**). Semi-quantitative intensity scoring revealed compartment- and response-specific patterns (**Suppl. Table S4**). Within the TA-stroma, IgG deposition was uniformly moderate-to-high across groups, whereas IgA expression was diminished in non-responders (**Suppl. Fig. S9O**). Moderate-to-high IgA expression was observed within the TE of both groups (**Fig. 6F, G**), but IgG intensity trended to be higher in post-BCG TIME from non-responders relative to that of pre-treatment tumors (**Fig. 6F, G**). These findings indicate that IgG-skewed antibody responses in non-responders are reflected at both systemic and tissue levels.

### Repeated intravesical BCG administration induces systemic immunosuppressive cytokine shifts in non-responders

To determine the cytokine profiles related to those of altered B cells, proteomic profiling of 96 plasma proteins across the three timepoints was performed (n = 25 patients; **Fig. 6A**, **Suppl. Table S1/Cohort 1**). Significantly increased levels of over 40 plasma proteins were observed post 4^th^-BCG instillation in all patients (**Fig. 6H**; **Suppl. Fig. S9P, Q and S10A**). BCG therapy triggered a pronounced Th1 cytokine response (IL-2, IL-12, IL-15, IFN-γ, TNF), along with pro-inflammatory mediators (IL-6, IL-18, TWEAK) and immune cell recruiting chemokines (CXCL9, CXCL10, CCL19, **Fig. H; Suppl. Fig. S10A**). Notably, the Differentially Expressed Proteins (DEPs) exhibited substantial overlap between post-1^st^ and post-4^th^ BCG groups (**Fig. 6I**), with increased levels of immune checkpoint proteins (PD-L1, PD-L2, and LAG-3). The sustained elevation of TNF and its cognate receptors (TNFRSF12A, TNFRSF21, TNFRSF4, TNFRSF9) further indicated persistent inflammatory signaling and prolonged immune activation (**Fig. 6H; Suppl. Fig. S10A**).

Elevated expression of stress-associated proteins such as MICA/B, HO1, and HGF was observed following post-4^th^ BCG instillation in all patients (**Fig. 6H, I**). Comparative analysis between responders and non-responders further revealed elevated levels of ADA, KLRD1, and GZMH in responders, while non-responders showed consistent expression of VEGFA at both pre-BCG and post-4^th^ BCG timepoints (**Fig. 6J**; **Suppl. Fig. S10B**). Temporal comparisons across treatment timepoints revealed dynamic shifts in circulating protein levels, with non-responders exhibiting a significant increase in IL-10, TNF, and TNFRSF21 levels post-BCG (**Fig. 6K; Suppl. Fig. S10C, D**).

Pathway enrichment analyses (Gene Ontology (GO) and Kyoto Encyclopedia of Genes and Genomes (KEGG)) corroborated these findings, highlighting significant involvement of immune activation pathways, including positive regulation of T cell activation, leukocyte migration, leukocyte proliferation, cytokine-cytokine receptor interaction, and TLR signaling (**Suppl. Fig. S10E, F**). Furthermore, spearman correlation analysis revealed significant positive correlations between IFN-γ and the chemokines CXCL9 and CXCL10 (**Suppl. Fig. S10G, H**), highlighting a Type I immune response following repeated mucosal exposure to BCG. Furthermore, the majority of DEPs across pre- and post-BCG timepoints (post-1^st^ BCG, post-4^th^ BCG) exhibited coordinated positive correlations, irrespective of clinical response status. These findings indicate that intravesical BCG treatment elicits a progressive, systemic immune activation characterized by T cell activation, pro-inflammatory signaling, and co-stimulatory engagement.

GO enrichment analysis further substantiated this immunosuppressive shift in non-responders, revealing pathways involved in negative regulation of B cell proliferation, chronic inflammation, and altered IFN responses (**Fig. 6L**). Spearman correlation analysis revealed significant negative associations between the cytotoxic T cell marker CD8A, IFN-γ and CXCL9 at the two timepoints in non-responders (**Suppl. Fig. S10G**). Such associations were absent in responders (**Suppl. Fig. S10H**). Notably, in non-responders, CD8A levels showed a positive correlation exclusively with the chemokine CCL23 (**Suppl. Fig. S10G**). The positive correlation between CCL23 with PD-L1 reinforces its association with pro-inflammatory yet immunosuppressive signaling.

## Discussion

Intravesical BCG-induced local and systemic immune shifts have largely been independently explored in patients with high-risk NMIBC despite emerging evidence on the significant influence of their coordinated interactions dictating the downstream anti-tumor mucosal immunity in the bladder microenvironment. Given that B cells are sentinels of mucosal immune system, in this study, we applied a unique approach to characterize B cell associated local and systemic responses in patients with high-risk NMIBC undergoing induction BCG therapy. Our novel findings identified B cell exhaustion and expansion of an ABC-biased humoral immune program, as a previously unrecognized determinant of poor response to BCG immunotherapy. Application of a holistic approach revealed that within the systemic and tumor local immune compartments, non-responders exhibit enrichment of ABCs and disruption of TA-TLS architecture. These unique features position dysfunctional mucosal immunity as a central feature of early tumor recurrence following adequate BCG therapy. Our findings suggest that a chronically altered pre-treatment inflammatory microenvironment potentially amplifies the pro-inflammatory cytokine milieu, predisposing the immune system toward ABC expansion even before BCG exposure[9]. Repeated antigenic stimulation via BCG antigens in a pre-existing systemically dysregulated immune state could therefore further promote immune exhaustion, limiting the adaptive anti-tumor immune capacity required for durable response to BCG immunotherapy.

Spatial analyses of tumors revealed a correlation between systemic ABC expansion and dysfunctional intratumoral immune architecture. In responders, TLS were enriched for DCs and cytotoxic T cells, with ABCs positioned at the TLS periphery - a configuration compatible with GC activity and coordinated T and B cell interactions. In contrast, non-responders exhibited accumulation of ABCs within TA-TLS cores and the TE. In the TA-stromal region, the presence of ABCs may thus contribute to local immunosuppression and impaired CD8^+^ T cell effector function, creating an environment permissive for tumor progression. The TIME of non-responders also showed an increased density of T_regs_ and CD163⁺ macrophages, together with transcriptional signatures of EF B cell activation and plasma cell differentiation. CD163⁺ macrophages secrete IL-10, TGF-β, and VEGF, factors known to suppress GC formation and promote tolerogenic B cell programs, while T_regs_ deplete IL-2 and inhibit effector T cell and DC activity within the TIME[46–50]. These observations align with emerging data across cancers that mature, GC–rich, TLSs predict treatment responses and favorable outcomes, whereas extrafollicular or exhausted B cell niches correlate with poor outcome[51–53].

Multiplex spatial immunophenotyping-based neighborhood analysis revealed that ABCs, T_regs_, and memory cytotoxic T cells were positioned in close proximity to the TE in non-responders. In responders, ABCs were integrated within domains enriched for helper T cells and myeloid populations. In tumors from non-responders, ABC-containing domains were enriched for cytotoxic T cell subsets and lacked myeloid and helper T cell co-enrichment, suggesting the emergence of spatially segregated B cell dominant niches lacking the antigen-presenting infrastructure required for effective adaptive priming, whereas in responders, ABCs and memory T cell populations were spatially excluded from the epithelial front, while effector T cells maintained epithelial proximity. Spatial transcriptomics corroborated these structural changes at the molecular level. Non-responders showed enrichment of genes associated with extrafollicular plasma cell differentiation, whereas responders displayed increased expression of genes associated with antigen presentation and DC recruitment. These findings suggest an immunosuppressed tumor epithelial niche in non-responders.

Among the molecular signatures of ABCs within the bladder TIME, *FCRL5* emerged as the most consistent and abundant marker. FCRL5 expression was observed in circulating ABC populations, highly expressed in spatial transcriptomic clusters, and associated with both reduced RFS and PFS in two independent cohorts of a total 409 patients. Experimental work in mouse models has shown that enforced *Fcrl5* expression breaks B cell anergy, enhances TLR responsiveness, and triggers systemic autoimmunity, establishing FCRL5 as an active driver of tolerance breach rather than a passive exhaustion marker[54, 55]. Another novel finding in our study was the presence of IFN-γ–expressing ABCs detected in most non-responder tumors, consistent with features of chronic, non-resolving inflammation. This finding aligns with a potential role of ABCs in IFN-γ induced adaptive immune resistance as a critical factor underlying poor response to BCG[56].

Peripheral ABC frequencies were elevated prior to treatment in non-responders and expanded further with repeated BCG instillations, indicating a persistent host immune configuration rather than a transient response to BCG. Single-cell transcriptomic analysis of circulating B cells confirmed enrichment of canonical ABC-associated transcriptional signatures, including *TBX21, FCRL5, ZEB2,* and *ITGAX*, before therapy initiation, which persisted following treatment. These findings align with emerging evidence that atypical or exhausted-like B cell states accumulate under chronic antigenic stimulation and IFN-rich environments and exhibit impaired recall capacity despite transcriptional activation[26, 32, 57, 58]. These observations are suggestive of a pre-existing ABC-biased immune architecture that may limit productive B cell maturation under strong mucosal immune stimulation during repeated BCG instillations.

Among factors that promote ABC differentiation, older age, carcinogen-induced chronic mucosal inflammation, and immune activation associated with repeated BCG administration are all constitutive features within majority of patients with high-risk NMIBC[35, 41]. Consistent with this, plasma proteomic profiling in non-responders revealed enrichment of IFN- and cytokine-driven inflammatory pathways associated with chronic B cell stimulation. Repeated BCG exposure may therefore reinforce an exhausted TIME rather than generate effective humoral immunity in those who experience early recurrence. Indeed, in our prior report based on an aging murine model of carcinogen-induced bladder cancer, repeated BCG instillations drove systemic and local ABC expansion and induced B cell-dependent changes in bone marrow composition, providing a pre-clinical basis for the patterns that we observed in human NMIBC[9]. Aligning with these findings, studies in both BCG vaccination and bladder cancer contexts show that BCG reprograms hematopoietic progenitors within the bone marrow, altering the composition and function of newly generated immune cells[21, 59–61]. While our cohorts included patients between 29-89 years, more than 70% patients were over 65 years old. Our finding on variable ABC frequencies in responders potentially also reflects the influence of older age, biological/chronological aging associated immune dysfunction or chronic comorbidities.

Our finding on post-BCG naïve B cell expansion in the responders in contrast to memory B cell expansion in non-responders, is supported by mechanistic evidence from our pre-clinical investigations[9], suggesting that effective BCG therapy requires recruitment of naïve, antigen-inexperienced B cells capable of mounting effective humoral responses. Loss of unswitched memory cells - normally a reservoir of clonal diversity - suggests progressive skewing of the B cell compartment toward terminal or dysfunctional states. This pattern mirrors observations in chronic inflammatory diseases where ABCs arise from memory B cells, display reduced BCR signaling, and preferentially engage innate-like or extrafollicular differentiation programs[25, 31, 32].

Serological profiling reinforced this shift toward extrafollicular humoral responses. Non-responders exhibited reactivity to significantly higher autoantigens between pre- and post-BCG timepoints, compared to responders. A recent pan-cancer B cell atlas encompassing 477 specimens across 20 tumor types established that tumors dominated by extrafollicular B cell programs are associated with inferior clinical outcomes compared with GC–driven immune responses[24, 27]. Consistent with this framework, the antibody repertoire in non-responders was characterized by *IGHG1* dominance, broad autoreactivity, and progressive anti-BCG IgG accumulation. Unlike GC-derived antibodies, extrafollicular B cell derived antibodies undergo limited affinity maturation. Thus, the expanding BCG-specific IgG response observed in non-responders likely represents immunological misdirection of antigen engagement without protective mucosal immunity, in concordance with previous reports[62]. Transcriptional evidence of IgG-skewed humoral output was also reflected within the TIME. IgG deposition in TE increased following BCG in non-responders, whereas IgA - a key mediator of mucosal immune homeostasis - was reduced within TA-stroma. These findings suggest that B cell responses in non-responders shift from mucosal effector programs toward systemic IgG-dominated responses that may be poorly suited for local tumor control.

Findings from this study carry important clinical implications. The stability of peripheral ABC frequencies across treatment timepoints suggests that systemic ABC profiling could serve as a minimally invasive biomarker to identify patients at high risk of BCG failure prior to therapy initiation. ABCs in our cohort consistently expressed PD-1 and PD-L1 and resided within immune checkpoint-rich niches, providing a strong rationale for combining BCG with immune checkpoint blockade, specifically, in patients with increased ABCs within their treatment naïve TIME. This concept is supported by recent clinical data demonstrating improved outcomes in a subset of patients treated with combination anti-PD-1 and BCG immunotherapy in the first line treatment setting of high-risk NMIBC. This phase 3 CREST trial demonstrated that sasanlimab, a subcutaneously administered anti-PD-1 monoclonal antibody, in combination with BCG induction and maintenance, reduced the risk of disease-related events by 32% compared with BCG alone in BCG-naïve high-risk NMIBC[63]. A modest survival benefit imparted by immune checkpoint inhibitors[64] further emphasize the need to precisely define the local and systemic immune features reflecting pre-treatment global mucosal immune exhaustion. While correlative biomarker profiles are awaited, findings from our study will also have significant implications in the immunomodulatory effects by other novel therapeutics such as Cretostimogene grenadenorepvec, Gemcitabine-Docetaxel chemotherapy and IL-15 receptor agonist that have shown favorable outcomes in BCG naïve patients.

This study is not without limitations. The cohort size, while sufficient to identify consistent immunological trends across multiple complementary modalities, limits the ability to perform a sex-disaggregated analyses. Female sex is an established risk factor for more aggressive NMIBC, and our prior murine model-based work showed a sex-differential pattern of B cell/ABC response to BCG. While abundance of memory B cells reflected lower pre-existing BCR diversity, further characterization of BCG response associated local and systemic BCR repertoire will inform the productive rearrangements within this compartment. Lastly, we did not incorporate the elements of the complement pathway in this study despite their emerging significance in the TA-TLS functions. Such studies will be essential to determine whether targeting ABC differentiation or function can restore productive humoral immunity during BCG therapy.

In summary, our findings indicate that a pre-existing ABC-biased immune state may constrain the capacity of BCG to elicit effective humoral immunity in NMIBC. In non-responders, and patients with depleted naïve B cell pools and reduced differentiation capacity, repeated BCG exposure is associated with amplification of systemic inflammatory signals and persistence of an FCRL5^+^ ABC-dominant B cell compartment, accompanied by reduced frequencies of unswitched memory B cells and expansion of autoreactive IgG responses. Within the bladder microenvironment, ABCs accumulate within TLS cores and IFN-enriched basal epithelial niches, potentially disrupting GC organization and promoting tumor permissive immune microenvironment. Recognition of this ABC-dominated immune architecture highlights an opportunity for predictive biomarker-guided therapeutic strategies in bladder cancer.

## Acknowledgements

This study was supported by research operating grants from the Canadian Institutes of Health Research, Bladder Cancer Canada, Leo and Anne Albert Institute for Bladder Cancer Care and Research, Cancer Research Society, and Canada Foundation for Innovation. We thank Lab Research Services at the Department of Pathology and Molecular Medicine, KHSC, for assistance with sectioning of archival FFPE tumor tissues. We thank QLMP for assistance with slide scanning on Halo software and Katy Milne at BC Cancer’s Molecular and Cellular Immunology Core (MCIC) for immunofluorescence staining. We thank Neil Winegarden (10X Genomics) for support with establishing scRNA-seq protocol for B cells.

## Author contributions

MK conceptualized and designed the study. PY, KS and GC performed experiments included in this study, analyzed data and contributed to manuscript writing and reviewing. AB and AG consented and collected blood from the KHSC patients. SP performed UBC sample processing under supervision of MR and PB. AA helped with reviewing histopathological features and retrieving of the archival tumor tissue specimens under the supervision of DMB. MJ contributed to writing, reviewing, editing of the manuscript. DRS and DMB helped with clinical classifications, study design and contributed to patient recruitment and selection. CG and MA optimized and conducted mIF experiments guided by SVR. HY and AH contributed to plasma Olink profiling and analysis. DC guided scRNA-seq analysis. RL contributed to study design and interpretation. TS and SL generated and analysed the transcriptome profiles for independent validation under supervision of LD. All authors reviewed the manuscript.

## Methods

### Human Ethics and Patient Cohort

This study was conducted in compliance with institutional ethical guidelines following approval by the Health Sciences Research Ethics Review Board at Kingston Health Sciences Centre (KHSC), Queen’s University, and University of British Columbia’s Clinical Research Ethics board. The study is in compliance with ethical standards set out by the Canadian Tri-Council Policy Statement: Ethical Conduct for Research Involving Humans-TCPS2, Canada. Written informed consent was obtained from all participants.

### Histopathological Evaluation

Hematoxylin phloxine saffron (HPS) - stained whole tumor sections from TURBT specimens (pre-BCG and post recurrence) were accessed from the KHSC pathology tissue archives (**Suppl. Table S1/Cohorts 1 and 2**). HPS-stained slides from 2-4 tumor tissue blocks per patient were scanned at high-resolution (20X) using the Olympus VS120 scanner. Digital images were subsequently uploaded to the web-based HALO image hosting platform by the QLMP histology core, facilitating digital access and annotation.

Histopathological evaluation of the tumor sections was conducted to assess the tumor stage, grade, invasion, immune cell infiltration and distribution patterns. Only the tumor regions with intact urothelium-defined as the presence of preserved, continuous, non-ulcerated bladder epithelium were included in downstream analyses. In the context of NMIBC, retention of an intact urothelial layer often denotes early-stage (Ta/T1) disease where CIS or papillary lesions remain confined to the mucosa or submucosa[65]. These regions are crucial for analyzing immune infiltration patterns at the epithelial–stromal interface, which is central to mucosal immune surveillance and BCG-induced immune activation.

Slides for spatial immunophenotyping were selected based on several histological criteria (**Fig. 1A**): (i) presence of TE with adjacent stroma, (ii) evidence of inflammatory infiltrates, particularly lymphoid aggregates within the LP or peri-tumoral immune cell clustering in the form of TLSs, (iii) preservation of urothelial architecture without significant crush artifact or cautery distortion, and (iv) minimal necrotic or hemorrhagic areas that could compromise antigen preservation. Serial 4 µm sections of selected formalin fixed paraffin embedded (FFPE) tissue blocks was performed by the KHSC histology core for mIF staining.

### Spatial immunophenotyping on whole tumor sections using multiplex immunofluorescence

Spatial immunophenotyping was conducted on 31 whole FFPE tumor specimens (18 BCG responders, and 9 BCG non-responders; pre- and post-recurrence). Two out of four serial sections per FFPE block were subjected to automated mIF staining at the Molecular and Cellular Immunology Core facility (Deeley Research Centre, Canada) as per our previously reported methods. Tissue sections were profiled using two eight-plex panels - the ABC panel and the T cell exhaustion (T_ex_) panel - each comprising fluorophore-conjugated primary antibodies directed against cell-surface epitopes to enable simultaneous spatial resolution of distinct immune cell populations (**Suppl. Table S2**). Stained sections were digitized using the Vectra Polaris multispectral imaging platform (Akoya Biosciences, USA), reviewed in PhenoChart software (v1.1.0), and subsequently imported into inForm Viewer (v2.5.0) for manual region of interest (ROI) annotation and high-magnification tile capture.

ROIs were systematically classified into four discrete categories based on the presence of mature or immature tumor-associated tertiary lymphoid structures (TA-TLS), immune cell infiltration within the tumor-associated stroma (TA-stroma), and lamina propria (LP) compartments (**Fig. 1C; Suppl. Fig. S1A**). Tissue sections yielding fewer than four ROIs per category were excluded from downstream analyses. Mature TA-TLS were defined as organized lymphoid structures situated within 1 mm distance from the TE, harboring a morphologically distinct GC with a minimum cellularity threshold of >1,000 DAPI⁺ nuclei (**Fig. 1A**). Immature TA-TLS were defined as nascent lymphoid aggregates devoid of noticeable GC architecture, with intermediate cellularity ranging from 100 to 1,000 DAPI⁺ nuclei.

To further enable high-plex spatially resolved phenotyping and quantitative cellular neighborhood analysis within intact tissue architecture, a representative subset of 12 slides (**Suppl. Table S1/Cohort 1**) was further interrogated using a 25-plex antibody panel identifying T, B and myeloid cell subsets (**Suppl. Table S2**) cyclic immunofluorescence approach on the PhenoCycler-Fusion platform (Akoya Biosciences), which couples the PhenoCycler fluidics instrument with the Fusion high-resolution fluorescence microscope to facilitate sequential, iterative antibody-based staining and imaging. Tissue staining and image acquisition were performed at the Division of Experimental Medicine, McGill University, in strict accordance with the facility’s validated standard operating procedures and manufacturer’s specifications[66]. To preserve the spatial integrity and cellular heterogeneity of the TIME, two histologically intact ‘tumor islands’ per sample meeting the above four listed criteria were selected for downstream analysis rather than discrete ROI sampling (**Fig. 3A**).

Downstream image processing and quantitative analysis across all three panels were performed using QuPath (v0.5.1; https://qupath.github.io). Single-cell segmentation was executed using StarDist, a convolutional neural network-based extension for QuPath, trained on fluorescence microscopy images for robust nuclear detection across tissue regions [66, 67]. Cell phenotype classification thresholds were established through supervised training on duplicate-channel images with manual boundary curation, and threshold accuracy was orthogonally validated against DAPI nuclear counterstaining. Co-expression of lineage and functional markers was resolved using the composite object classifier framework within QuPath. For ABC and T_ex_ panel data, per-cell quantitative metrics, including cell densities and spatial coordinates, were exported to GraphPad Prism (v10.0) and R (v4.3) for group-wise statistical comparisons, with per-section ROI means as the unit of analysis.

For the PhenoCycler-Fusion dataset, single-cell x,y spatial centroid coordinates and per-marker mean fluorescence intensities (MFI) were exported as structured .csv files and processed in R for cell-cell proximity analyses, spatial interaction modeling, and cellular neighborhood quantification.

### Spatial neighborhood analysis

Spatial proximity between cell populations was quantified using k-nearest neighbor distance analysis (k-distance)[67]. Specifically, distances were computed from all queried cell types to two reference cell populations: ABCs and tumor epithelial cells. For each cell, the mean distance to its 10 nearest neighbors (k = 10) belonging to the reference population was calculated[67]. The distribution of distances from immune cells to cancer cells was visualized using ridge density plots, stratified by ROI and treatment response category. Ridge plots were generated using the R packages ggridges (0.5.6) and ggplot2 (4.0.1) as defined in previous reports[67]. To statistically compare spatial proximity across treatment response groups, violin plots with embedded boxplots were generated. The mean value for each group was overlaid, and differences between groups were assessed using the Kruskal–Wallis test.

### Cellular distance and Niche analysis

To quantify spatial relationships between different cellular phenotypes, minimum distances between all pairs of cell types were calculated using *calculate_minimum_distances_between_celltypes()* function in SPIAT (Spatial Image Analysis of Tissues) [v1.10.0] R package. For each pair of cell phenotypes, the minimum distance (Nearest neighbour) between cells was computed and summarized using *calculate_summary_distances_between_celltypes()* function to generate mean interaction distances which was later visualised as heatmaps using pheatmap package (1.0.13). To infer spatial cellular niches, we used spatial topic (SpaTopic) modeling-based spatial clustering approach[68]. The model was run with eight spatial topics (ntopics = 8), a Gaussian spatial smoothing parameter (σ) of 50, and a neighborhood radius of 400 µm to capture local cellular context using function *SpaTopic_inference().* This approach infers latent spatial domains that represent recurring cellular neighborhood patterns within the tissue microenvironment. Posterior topic probabilities were computed for each cell, and cells were assigned to the spatial domain corresponding to the highest posterior probability. To characterize the phenotypic composition of each inferred spatial domain, topic-marker association matrices were generated and visualized using hierarchical clustering heatmaps, enabling identification of cell type enriched spatial niches. To visualize the cellular composition of the inferred spatial niches, the number and relative proportion of each cell phenotype within each spatial topic were calculated using data manipulation functions from dplyr (1.1.4) and tidyverse (2.0.0). These summaries were visualized using dot plots generated with ggplot2 and styling utilities from ggpubr (0.6.0). Further cells expressing PD-1, PD-L1 and IFN-γ were plotted using *circos.trackPlotRegion()* function from circlize package (0.4.16).

### Preparation of sample for spatial transcriptomic profiling using Xenium In Situ platform

FFPE bladder tumor whole sections from six patients first characterized using 25-plex mIF described above (**Suppl. Table S1/Cohorts 1 and 2)** were used to select areas for single cell spatial transcriptomic profiling using Xenium In Situ platform with the Prime 5K Gene Expression Panel (10X Genomics). Tissue sectioning (5 µm thickness), slide preparation, and Xenium In Situ processing was performed by the Pathology Research Program Laboratory at the University Health Network (Canada), strictly following the manufacturer’s recommended workflow and quality control guidelines (CG000578, 10X Genomics). Two Xenium slides were generated, each containing three tissue sections within the 12 mm x 24 mm imageable area.

The tissue areas were selected based on histopathological review of H&E-stained sections and mIF-based spatial immune profiles to ensure inclusion of intact TE and TA-TLS. Following Xenium In Situ imaging and transcript detection, slides underwent post-assay H&E staining for morphological validation at the PMGC. Spatial transcriptomic profiling was performed using approximately 5,000 targeted RNA probes. The resulting raw data outputs, including the feature-cell count matrix (HDF5 format), transcript-level localization files, cell boundary annotations (CSV format), and the experiment.xenium configuration file for visualization in Xenium Explorer were generated by the processing facility and transferred to our group via a secure SFTP server. All downstream analyses described in this study were conducted exclusively using these raw Xenium output files.

### Spatial transcriptomics data processing and clustering

Spatial transcriptomics data generated using the Xenium platform were processed using the Seurat (v5) framework in R. Quality control filtering was performed to remove low-quality cells based on transcript counts. The distribution of detected transcripts per cell was visualized using a histogram, and a lower threshold of 20 transcripts per cell was selected as the cutoff for inclusion. Cells with transcript counts above the 98^th^ percentile were also excluded to remove potential outliers as mentioned in protocol provided by 10X Genomics data analysis guide (https://github.com/10XGenomics/analysis_guides/blob/main/Xenium_5k_data_analysis_journey.ipynb). The filtered dataset was normalized using median-based scaling implemented in the *NormalizeData()* function, with the scale factor set to the median number of transcripts per cell. Highly variable genes were identified using *FindVariableFeatures()* function with default parameters. For dimensionality reduction, the Xenium assay was set as the default, and the data were scaled using *ScaleData()*. PCA was then performed using *RunPCA()* function, and the number of informative principal components was determined by inspection of an elbow plot. To correct for potential batch effects across samples, data integration was performed using the Harmony algorithm via the *IntegrateLayers()* function, generating a batch-corrected low-dimensional embedding. UMAP was subsequently applied to the Harmony embedding using *RunUMAP()* function and visualized using ggplot2 package.

Cell neighbors were constructed using *FindNeighbors()* function, followed by unsupervised clustering using the Louvain clustering algorithm implemented in *FindClusters()* function with a resolution parameter of 0.12. To enable downstream differential expression analysis, assay layers were joined using *JoinLayers()* functions. Cluster-specific marker genes were identified using *FindAllMarkers()* function, retaining only genes with positive differential expression and was manually annotated using conventional phenotypic markers (**Suppl. Table S5**). Further, heatmap was plotted using SeuratExtend package (1.2.7) where z-score was calculated for each cluster using *CalcStats()* function. Gene set analysis was performed using *GeneSetAnalysis()* function in SeuratExtend package for Hallmark, Reactome and GO pathways and results were later plotted as heatmaps using *Heatmap()* function.

### Subset clustering and pathway analysis of spatial transcriptomics data

To investigate transcriptional heterogeneity within B cell and epithelial-lineage populations, cells annotated as B and plasma cells, and epithelial cells were subset separately from the integrated dataset generated using *subset()* function in Seurat. As described above dimensionality reduction was performed with 15 dimensions and resolution of 0.2 and z-score scaled marker gene statistics were calculated and visualized using heatmaps (**Suppl. Table S5**). Clusters were subsequently labeled as B cell subclusters and ordered numerically for downstream analyses. Gene sets corresponding to naïve B cells, memory B cells, atypical B cells, GC B cells, proliferating cells, and plasma cells were compiled based on previously reported lineage marker genes[23]. Epithelial markers were selected based on previously published papers[69–71]. Expression of these gene signatures across clusters was visualized using dot plots to highlight subtype-specific transcriptional programs. To further characterize functional differences between populations across treatment conditions, gene set enrichment analyses were performed using pathway collections from MSigDB, Hallmark gene sets and Reactome. Differential pathway activity across clinical groups defined by BCG treatment response timepoints pre-BCG responder, pre-BCG non-responder, and post-BCG non-responder) was visualized using heatmaps. UMAP visualizations were additionally generated to illustrate the distribution of B cell subclusters across patient samples and clinical response groups.

### Spatial neighborhood and cellular niche analysis for spatial transcriptomics

To characterize spatial cellular niches within the tissue microenvironment, neighborhood-based analysis was performed on spatial transcriptomics data processed with Seurat in R (<u>Chatomics</u>). Cells from each sample were extracted individually from the integrated dataset for spatial niche analysis. Spatial coordinates corresponding to cell centroids were retrieved from the Xenium image data and used to compute spatial neighborhoods. A fixed-radius nearest neighbor search was implemented using the dbscan R package (1.2.2), where neighbors within a radius of 50 µm were identified for each cell. The resulting neighborhood graph was used to quantify the composition of surrounding cell types for every cell by counting the frequency of each annotated cell type within its local neighborhood. These neighborhood composition vectors were assembled into a cell-by-cell matrix representing the abundance of each cell type in the local spatial context. To identify recurring spatial patterns, the matrix was subjected to k-means clustering with k = 13, grouping cells into distinct spatial niche categories based on their surrounding cellular composition. For additional clustering and visualization, the neighborhood composition matrix was converted into a Seurat object and normalized using SCTransform. Dimensionality reduction was performed using principal component analysis, followed by construction of a nearest neighbor graph and unsupervised clustering using the Louvain clustering algorithm. The resulting niche clusters were visualized using UMAP.

To further characterize spatial niches, a dedicated niche assay was generated using the *BuildNicheAssay()* function with 30 nearest neighbors per cell and five niche clusters. Clustering concordance between k-means and graph-based approaches was evaluated using pairwise Jaccard similarity heatmaps, implemented using the scclusteval (0.0.0.9000) R package. The average abundance of surrounding cell types within each spatial niche cluster was calculated and visualized using heatmaps generated with ComplexHeatmap (2.24.0). Additional dot plot visualizations were produced to summarize cell-type distributions across niche clusters. Finally, spatial distributions of niche clusters were projected back onto the original tissue coordinates using the *ImageDimPlot()* function, enabling visualization of spatially organized cellular neighborhoods within the tissue section. Cluster assignments for cell types, B-cell and epithelial cell subclusters, and spatial niche clusters were exported as tabular metadata files for downstream visualization in Xenium explorer.

### Isolation of B cells from peripheral blood

Peripheral blood samples were prospectively collected from patients diagnosed with high-risk NMIBC receiving BCG immunotherapy at the Hotel Dieu Hospital, KHSC (cohort 1 and 2), and Vancouver Prostate Center (cohort 3), Canada. Peripheral blood (15 – 20 ml) was collected at three timepoints during BCG induction phase (**Fig. 5A**): prior to treatment (pre-BCG, n = 31), following the first dose (post-1^st^ BCG, n = 26), and after the fourth dose (post-4^th^ BCG, n = 29). Only samples from patients who received an adequate dose of BCG (5 induction + 2 maintenance doses), as per established guidelines, were included in the study[4, 5, 72]. Clinical information, including patient demographics, tumor grade, stage, and treatment outcomes are summarized in **Suppl. Table S1/Cohorts 1, 2 and 3**. Patient confidentiality was maintained throughout the study, and all clinical data were anonymized prior to analysis.

Peripheral blood (15 – 20 ml) was collected in ethylenediaminetetraacetic acid (EDTA)-coated tubes by venipuncture. All samples were processed within 30 – 60 mins following collection. Blood samples were centrifuged at 800 x g (brakes off) for 10 mins at 4℃ to separate the buffy coat from plasma and red blood cell (RBC) layer. Plasma was aliquoted and stored at - 80℃. The buffy coat was subjected to total B cell isolation (Cohort 1 and 2; n = 31) using the magnetic bead based EasySep^TM^ Direct Human B Cell Isolation Kit (Cat# 19675RF, StemCell Technologies, Canada). All samples were processed on the RoboSep^TM^ – S (Program# 19674; Cat# 21000, StemCell Technologies) automated cell separation system (**Fig. 5A**).

### Profiling of circulating B cell subsets using multiparametric flow cytometry

Enriched B cells were resuspended in RoboSep Buffer (Cat# 20104, StemCell Technologies) and washed with 1X PBS. A total of 0.25 - 0.5 × 10^6^ cells/well were seeded in a V-shaped 96-well plate and stained with a fixable viability dye (Cat#L34966, ThermoFisher, USA) and Human TruStain FcX^TM^ (Cat# 422302, BioLegend, USA) for 30 mins on ice. Subsequently, enriched B cells were stained using a custom designed panel of 7 fluorescent conjugated antibodies (BioLegend) to identify subsets of B cells including ABCs: CD19 (Alexa Fluor 700, Cat# 302226), CD21 (FITC, Cat# 354910), CD11c (Brilliant Violet 785, Cat# 301644), CD27 (PE/Dazzle 594, Cat# 356422), IgD (APC, Cat# 348222), IgM (Pacific Blue, Cat# 314514), and FCRL5 (PE, Cat# 340304) - for 30 mins at 4℃. Unstained controls, fluorescence minus one (FMO), and single-color controls were used for gating and subsequent analysis. The stained cells were then resuspended in 200 µl staining buffer (2% FBS in 1X PBS) and data were acquired on a CytoFlex-S Flow Cytometer (Beckman Coulter, Canada). The gating strategy for identifying ABCs and other B cell subsets is detailed in the **Suppl. Fig. S5A, B**. Flow cytometry data analysis was conducted using FlowJoⓇ software v10.10 (BD Biosciences, Canada). Calculated frequencies were exported to GraphPad Prism (v9.5.1, GraphPad Software, USA) for statistical analysis. Samples with more than 75% viability (**Suppl. Fig. S6A**) and CD19^+^ purity (**Suppl. Fig. S6B**) were used for downstream analysis. Four paired samples (pre-BCG and post-4^th^ BCG; **Suppl. Table S1/Cohort 2**) from both BCG responders and non-responders were used for concatenation for downstream analysis. Live CD19^+^ total B cells were gated, and 15,000 B cells per sample were down sampled using the DownsampleV3 plugin. B cell subsets were identified using the FlowSOM and visualized using tSNE plots in FlowJo.

To correlate systemic B cell profiles with the local immune microenvironment, FFPE tumor specimens from TURBT procedures at pre-BCG and post-recurrence were obtained from the KHSC pathology archives with the assistance of a genitourinary pathologists (AA and DM). In both the KHSC and UBC cohorts (**Suppl. Table S1/Cohorts 1, 2 and 3**), BCG non-responders (n = 15) were defined as patients exhibiting recurrence within 12 months of BCG treatment initiation, whereas patients who did not experience recurrence for >24 months were categorized as BCG-responders (n = 29).

### Sample preparation and fixation for scRNA-sequencing

B cells were isolated from four paired peripheral blood samples (Pre-BCG and Post 4^th^ BCG) using EasySep^TM^ Direct Human B Cell Isolation Kit, a magnetic bead-based enrichment procedures (Cat#19675RF, StemCell Technologies) and ultimately resuspended in PBS. Fixed single cell suspensions with more than 90% viability were prepared using Chromium Next GEM Single Cell Fixed RNA Sample Preparation Kit (Cat# PN-1000414, 10X Genomics, USA) as per manufacturer’s instructions and stored at −80℃.

Fixed B cell suspensions were then prepared for probe hybridization using the Chromium Fixed RNA Kit (Cat# PN-1000475, 10X Genomics). Whole-transcriptome probe pairs targeting human genes were added to each fixed sample and hybridized overnight, followed by a series of washes and quenching steps as detailed in the user guide. After hybridization, samples from each pair were multiplexed using the kit’s probe barcodes, pooled at equal cell numbers as allowed by the fixed RNA multiplexed workflow, and prepared for gel bead-in-emulsion (GEM) preparation.

### GEM generation, library construction, and sequencing

GEM generation and barcoding were carried out using a Chromium Next GEM Chip Q (Cat# PN-1000422, 10X Genomics) as per manufacturers protocol. The final libraries were size selected with SPRIselect beads and assessed on an Agilent Bioanalyzer High Sensitivity instrument at Queen’s Laboratory for Molecular Pathology (QLMP, Canada) prior to sending for sequencing. Pooled libraries were then sent to Princess Margaret Genomics Centre (PMGC, Canada).

### scRNA seq data processing and analysis

Following sequencing *Fastq* files were generated using the Cell Ranger Multi v8.0.1 (10X Genomics). Demultiplexed raw base call files (Feature/barcode matrix HDF5 raw) per sample was obtained after processing using the Cell Ranger count function, aligning to the human transcriptome (GRCh38) with default parameters – which generated gene-by-barcode count matrices for downstream analysis.

Gene expression libraries were imported into R (v4.5) and processed with Seurat v5 (https://github.com/satijalab/seurat/). Cells with more than 5% mitochondrial content were filtered out along with the following threshold: 300< nFeature ≤ 7,500 and 1,000 <nCounts ≤ 75,000). Cells were visualized using bar graph (**Suppl. Fig. S7A**) and violin plots after the preprocessing (**Suppl. Fig. S7B**). Filtered data were then normalized using *NormalizeData()* function of Seurat. The data were processed with principal-component analysis (PCA) and clustered using the Louvain algorithm (resolution = 0.25) on a nearest neighbor graph derived from first 30 principal components. Uniform manifold approximation and projection (UMAP) embeddings were generated using the *RunUMAP()* function on the first 30 principal components. *FindNeighbors(),* and *FindClusters()* function was used to cluster the cells. *FeaturePlot()* and *VlnPlot()* functions were used to visualize the clustered data. Using *presto::wilcoxauc(),* top 10 markers from each cluster was identified and used for manual annotation of the clusters. Only cells that were positive for B cell-associated genes (*CD19, MS4A1, CD79A, CD79B*) were included for downstream analysis (**Suppl. Fig. S7C**).

Pseudobulk differential expression was performed at the sample level by aggregating single-cell counts within each biological replicate and cell type, followed by bulk RNA-seq modeling with DESeq2. Pseudobulk count matrices for each B cell subset were analyzed independently using DESeq2 (v1.x) in R, modeling the negative binomial distribution of counts with gene-wise dispersion estimation and empirical Bayes shrinkage. Size factors were estimated by DESeq2’s median-of-ratios method to normalize for library size, and differential expression was tested using the Wald test with Benjamini–Hochberg adjustment of p-values to control the false discovery rate. Genes with an adjusted p-value < 0.05 and an absolute log2 fold change ≥ 0.25 were considered differentially expressed.

### Cell Communication Analysis

CellChat was applied to the Seurat object grouped by annotated B cell subsets, using the CellChatDB database for ligand-receptor pairs as mentioned in https://github.com/jinworks/CellChathttps://github.com/jinworks/CellChat. Cell-cell interactions expressed in < 10 cells were removed from downstream analysis and comparison. Overexpressed ligand and receptor genes were identified using the *identifyOverExpressedGenes()* function, followed by identification of enriched ligand–receptor interactions using *identifyOverExpressedInteractions()* function (**Suppl. Table S2**). Global communication networks were visualized using circular interaction plots, where edges represent the number or strength of predicted interactions between cell populations. Interaction probabilities were inferred for key pathways, with network centrality analysis (*netAnalysis_computeCentrality()* function) to identify sender/receiver roles among subsets. Cophenetic and Silhouette curves were plotted for incoming and outgoing signals to estimate the k value (k = 4 for incoming, k = 3 for outgoing) for the interaction analysis and visualized using alluvial plots (**Suppl. Fig. S8H, I**).

### Independent validation of FCRL5 expression in tumor bulk RNA-sequencing profiles

Normalized bulk RNA sequencing data from two independent cohorts were analyzed to evaluate the association between *FCRL5* expression and clinical outcomes post BCG treatment. The first cohort obtained from an unpublished dataset (n = 126) [*Strandgaard cohort*][43] was generated at Aarhus University, Denmark by Dyrsjkjot and colleagues. The second cohort of n = 283 patients [*de Jong cohort*] was extracted from previously published dataset^45^. Expression levels of *FCRL5*, a gene specifically expressed in ABCs, was used to stratify patients into three groups based on its distribution across all the samples in the cohort. Specifically, patients with *FCRL5* expression values ≤33^rd^ percentile were classified as “low,” those with values ≥ 66^th^ percentile as “high,” and the remaining patients as “intermediate.” High-grade RFS and PFS was assessed using Kaplan–Meier estimates. Survival curves were generated in R (v4.5.0) using the survival package (v3.8-3) and visualized with the survminer package (v0.5.1). Differences between groups were evaluated using the log-rank test.

### Circulating autoantibody profiling on HuProt™ human proteome microarray

Circulating levels of IgG autoantibodies were profiled using the ImmuneProfiler Assay on HuProt™ human proteome arrays (CDI Laboratories, Inc., USA), enabling high-throughput profiling of over 21,000 human proteins, including isoform variants and protein fragments (**Fig. 6A**)[73–75]. The raw sample-peptide matrices after checking for QC were imported into the R statistical environment (v.4.4.1) and processed using the limma microarray analysis suite (v.3.60.6) as described previously[73, 74]. Briefly, both microarrays and plasma samples were subjected to a 1-hr blocking step prior to the assay. Plasma samples were diluted 1:1,000 in CDISampleBuffer, while arrays were incubated with CDIArrayBuffer at room temperature (RT) under gentle shaking. Subsequently, each blocked and diluted sample was incubated on a HuProt™ microarray at RT for 1 hr with gentle shaking. Following the probing step, the arrays were washed three times for 10 min each with TBS-T (1x TBS containing 0.1% Tween 20). They were then incubated with Alexa Flour 647-conjugated anti-human IgG (Fc-specific) at RT for 1 hr in a light-proof box under gentle shaking. After incubation, the arrays were subjected to three additional 10 min washes with TBS-T, followed by three rinses with deionized water. Finally, the arrays were scanned using a GenePix 4000B scanner (Molecular Devices, USA) for data acquisition. Raw fluorescence intensity readings were collected from the scanner along with signal intensities of secondary antibody (Alexa 647 anti-human Ig) used in the study. The correlation coefficient (CC) for duplicate spots on each array, calculated as a part of the quality control process for IgG, averaged ∼0.97 (with a cutoff of 0.95). This indicated high reproducibility of duplicate spots and accurate grid alignment on the images, ensuring reliable data collection. The raw sample-peptide matrices were then imported into the R statistical environment (v.4.4.1) and processed using the limma microarray analysis suite (v.3.60.6) as described previously[74].

Background intensities for each sample were corrected using the *backgroundCorrect* function (method: normexp, normexp.method: rma). The data were then normalized across samples using *normalizeBetweenArrays* function (method: “cyclicloess”). For the filtering step, the median intensity of each peptide across all samples was calculated, and peptides exhibiting low reactivity or significant outlier values were excluded. Peptides with high median signal intensity in at least 3 samples were included for the analysis. Principal component analysis (PCA) was performed using the *prcomp* function (scale: TRUE), and the resulting loadings were visualized using the ggplot2 package.

To identify differentially abundant autoantibodies, a linear model was first fitted for each peptide to assess variation in peptide reactivity (against autoantibodies) across all arrays using limma’s *lmFit* function. Following model construction, a contrast matrix was generated to evaluate differences between treatment conditions (pre-BCG vs post-BCG) and treatment response groups (responder vs non-responder in pre-BCG and post-BCG condition). Differential abundance testing of autoantibodies was performed using the empirical Bayes method, as implemented in limma package. Resulting peptides were filtered based on an absolute log_2_(fold change) > 1 and a p value < 0.05. Differentially upregulated and downregulated autoantibodies (**Suppl. Table S4**) were visualized using volcano plots and boxplots, generated with the ggplot2 package.

Supervised and unsupervised clustering of the differentially abundant peptides was conducted, with the results visualized as heatmaps using the ComplexHeatmap package (v.2.20.0). GO enrichment analysis was conducted for autoantibodies showing increased abundance (log_2_ fold change > 1) using limma’s *goana* function. Biological process GO terms with a p value < 0.05 were selected and visualized using the ggplot2 package.

### Human Immunoglobulin Isotyping

Plasma Ig were analysed using the LEGENDplex^TM^ Human Immunoglobulin Isotyping panel (6-plex) by flow cytometry (Cat# 740640, BioLegend). Samples used in the assay are detailed in **Suppl. Table S1/Cohort 1,** Plasma samples and standards were prepared using assay buffer as per the manufacturer’s instructions. Data were analysed using the cloud-specific LEGENDplex Data Analysis Software Suite Qognit (v.2023.02.15, USA). Statistically significant differences in the levels of Ig isotypes between the groups were determined using GraphPad Prism (v.10.0.0).

### Detection of plasma anti-BCG antibodies

Lyophilized BCG-Tice (50 mg, Merck, Canada) was rehydrated in 1 ml PBS and incubated at RT for 1 hour. The bacterial suspension was centrifuged at 600 x g for 5 mins, and the pellet was resuspended in 1 ml lysis buffer consisting of 50 mM tris-hydrochloride, 0.5 mM EDTA, 60 mM sodium phosphate, 0.67% SDS, and protease inhibitor cocktail (Cat# 11836170001, Roche)[16, 76, 77]. Cells were lysed by sonication (15s pulses x 5 cycles). The lysate was sequentially centrifuged at 600 x g and 16,000 x g for 10 mins to remove insoluble debris. The supernatant was collected in a fresh tube and stored at −80℃ until further use. Protein concentrations were determined using the Pierce^TM^ BCA Assay (Cat# 23225, ThermoFisher).

BCG antigen whole lysates were diluted in 5X ELISA Coating buffer (Cat# 421701, BioLegend) as per manufacturer’s instructions and coated onto high-binding 96-well plate (Cat# 3590, Costar) at 100 ng/well. Plates were incubated overnight at 4℃. Wells were washed with PBS containing 0.05% Tween-20 (PBS-T) and blocked with 2% BSA in PBS-T for 2 hrs at RT. Following blocking, plates were washed with PBS-T and incubated with plasma samples diluted in blocking buffer (from 1:100 to 1:100,000; 10-fold dilution series). After washing, HRP-conjugated rabbit anti-human IgG H&L (Cat# ab6759, Abcam) was added and incubated for 1 hr at RT. Plates were washed with PBS-T, and 100 µl of TMB substrate Set (Cat# 421101, BioLegend) was added to each well and incubated for 15-20 mins at RT. The reaction was stopped using Stop Solution for TMB Substrate (Cat# 423001, BioLegend). Absorbance was measured at 450 nm with background correction at 570 nm using SoftMax Pro software (v7.1) on a microplate reader (SpectraMax^TM^ ABS Plus) within 30 mins of stopping the reaction (**Suppl. Table S3**).

### Plasma proteomic analysis on Olink platform

The levels of 92 proteins were measured in plasma samples collected at pre-BCG and post-4^th^ BCG time points, using the Olink® target Immuno-Oncology panel *via* multiplex proximity extension assay at the Icahn School of Medicine at Mount Sinai, USA (**Fig. 6A**). Results were reported as normalized protein expression (NPX) value, a log_2_ scale metric designed by Olink to facilitate relative protein quantification. The R package OlinkAnalyze (v.4.0.2) was used for the data analysis[78, 79]. To assess sample quality, *olink_qc_plot* and *olink_dist_plot* functions were used to generate a facet plot (interquartile range vs median for all samples, **Suppl. Fig. S9P**) and boxplots (NPX values vs sample ID, **Suppl. Fig. S9Q**), respectively. All samples passed QC and were included in subsequent analyses.

Paired comparisons across BCG treatment timepoints were performed using the non-parametric Wilcoxon signed-rank exact test was applied. For unpaired comparisons between responders and non-responders, the Wilcoxon rank-sum exact test was performed. Both analyses were conducted using the *olink_wilcox* function in OlinkAnalyze package. Temporal trends across treatment timepoints were assessed using ANOVA (olink_ordinalregression). DEPs were identified based on a p-value <0.05 and visualized with volcano plots (ggplot2 package, v.3.5.1). Spearman correlation evaluated relationships between samples and DEPs. and were visualized on heatmaps generated using the pheatmap package (v.1.0.12). Gene Ontology (GO) and Kyoto Encyclopedia of Genes and Genomes (KEGG) enrichment analyses for DEPs were conducted using the ClusterProfiler (v.4.12.6) and org.Hs.eg.db (v.3.19.1) packages, applying Benjamini–Hochberg correction method (adjusted p < 0.05). Significant pathways were visualized using dot plots (ggplot2).

### Statistical analysis

For statistical analyses performed using GraphPad Prism, results are expressed as mean ± SD. Comparisons between two groups with one independent variable was performed using non-parametric t-test (Mann-Whitney). Comparisons between groups with two independent variables were performed using one-way ANOVA with Tukey’s multiple comparison test. Differences among three groups were analyzed using non-parametric Kruskal-Wallis test with Dunn’s multiple test comparisons. Differences between two unpaired groups were analyzed using non-parametric multiple Mann-Whitney test with Holm-Šídák’s multiple comparisons test. Statistical significance is indicated by *p<0.05, **p<0.01, ***p<0.001, ****p<0.001, ns - not significant.

## Data availability

Data generated in this study are available in the manuscript and its Suppl. files. All materials and any other data are available from the corresponding author upon reasonable request. No custom codes were generated for the analysis (All the tutorials and standard pipelines have been mentioned in the protocol section).

