## Supplemental figures for "B cell exhaustion associates with poor response to Bacillus Calmette-Guérin immunotherapy in patients with bladder cancer"

### Supplementary Figures

***Figure S1. Spatial quantification of B cell subsets in the bladder tumor immune microenvironment.*** Schematic overview of the multiplex immunofluorescence (mIF) analysis workflow showing region-of-interest (ROI) selection across whole tumor sections and sampling across treatment groups, including responders (R) and non-responders (NR) at pre-BCG and post-4$^{th}$ BCG timepoints. Representative ROIs illustrate tumor-associated (TA) tertiary lymphoid structures (TA-TLS), TA-stroma, TA-lymphoid aggregates and lamina propria (LP) compartments analyzed across two staining panels (**A**). Box plots showing $log_2$-transformed cell counts of CD20$^+$ B cells, CD79a$^+$ B lineage cells, and CD79a$^+$CD20$^+$ cells across immature TA-TLS and LP regions stratified by response status (**B**). Bar plots summarizing the spatial distribution of CD20$^+$ B cells (**C**), CD20$^+$PD-1$^+$ cells (**D**), CD79a$^+$CD21$^{-/+}$CD27$^+$ cells (**E**) and CD20$^+$CD21$^{-/+}$CD27$^−$ cells (**F**) across tissue compartments including TA-stroma, mature TA-TLS, immature TA-TLS, and LP. Bar plots showing compartment-specific abundance of CD20$^+$CD21$^−$CD11c$^+$ (**G**), CD79a$^+$CD21$^−$CD11c$^+$PD-1$^+$ (**H**), and CD20$^+$CD21$^−$CD11c$^+$PD-1$^+$ (**I**) cells in responders and non-responders. Longitudinal comparisons of CD79a$^+$CD21$^−$CD11c$^+$ (**J**), CD79a$^+$CD21$^−$CD11c$^+$PD-1$^+$ (**K**), and CD20$^+$CD21$^−$CD11c$^+$PD-1$^+$ (**L**) cells across pre- and post-BCG samples. Stacked bar plots summarizing the proportional distribution of major immune cell populations across TA–immature TLS and LP regions, comparing responders with non-responders at pre- and post-BCG timepoints (**M**). Scatter bar plot showing spatial abundance of CD79a$^-$CD20$^−$CD11c$^+$ cells across tissue compartments stratified by clinical response (**N**). Each point represents an average of 3-6 ROIs per sample. Bars indicate mean ± SD. All statistical analyses were performed in GraphPad Prism using non-parametric tests; $*p < 0.05$, $**p <0.01$.

*Figure S1*

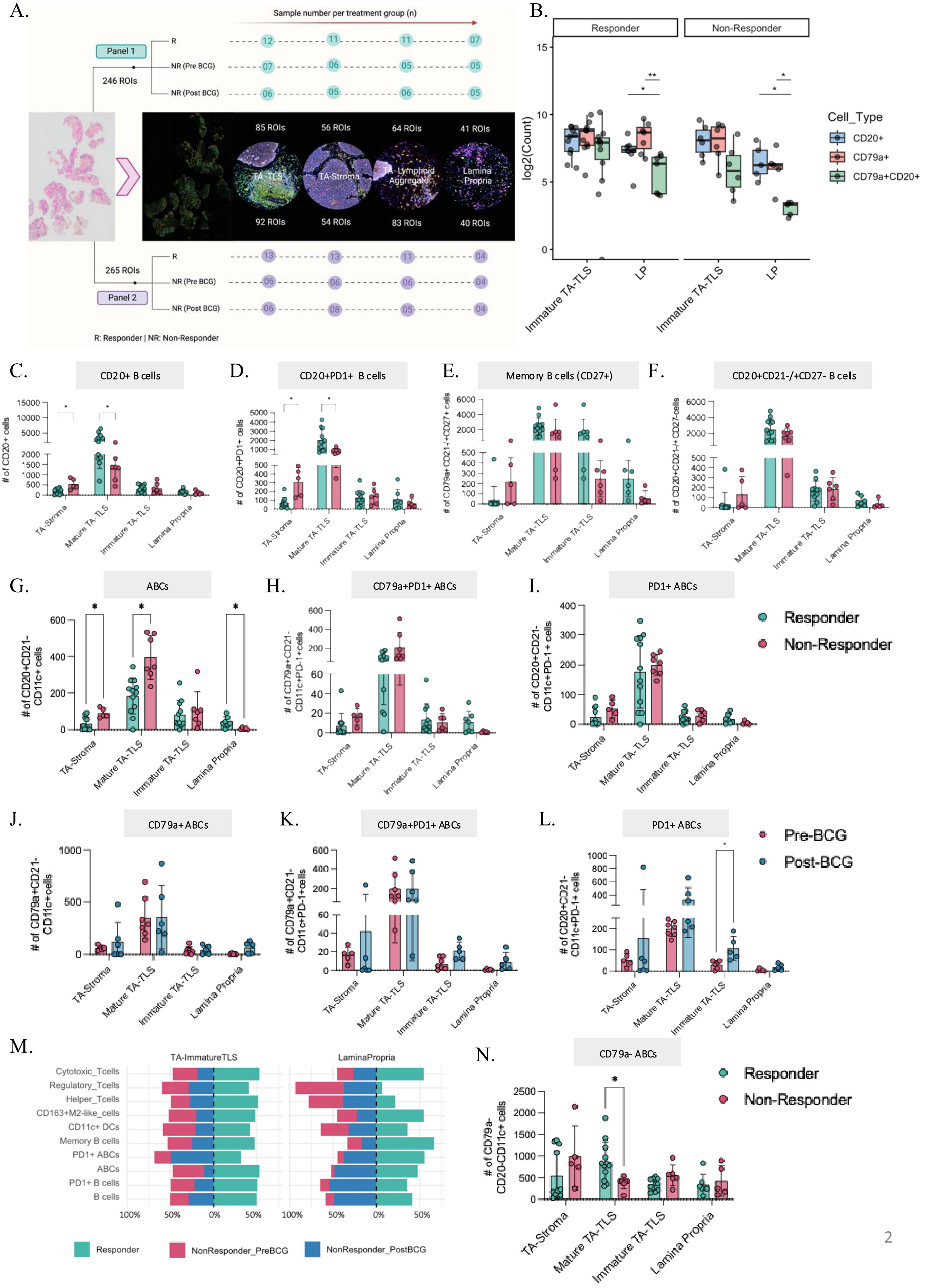
A.
Sample number per treatment group (n)
Panel 1
246 ROIs
R
NR (Pre BCG)
NR (Post BCG)
12 11 11 07
07 06 05 05
06 05 06 05
85 ROIs
56 ROIs
64 ROIs
41 ROIs
TA-TLS
TA-Stroma
TA- Lymphoid Aggregate
Lamina Propria
92 ROIs
54 ROIs
83 ROIs
40 ROIs
265 ROIs
Panel 2
13 13 11 04
06 06 06 04
06 08 05 04
R: Responder | NR: Non-Responder
B.
Responder
Non-Responder
log2(Count)
Immature TA-TLS
LP
Cell_Type
CD20+
CD79a+
CD79a+CD20+
C.
CD20+ B cells
### of CD20+ cells
D.
CD20+PD1+ B cells
### of CD20+PD1+ cells
E.
Memory B cells (CD27+)
### of CD79a+ CD21-/+CD27+ cells
F.
CD20+CD21-/+CD27- B cells
### of CD20+CD21-/+ CD27-cells
TA-Stroma
Mature TA-TLS
Immature TA-TLS
Lamina Propria
G.
ABCs
### of CD20+CD21- CD11c+ cells
H.
CD79a+PD1+ ABCs
### of CD79a+CD21- CD11c+PD-1+cells
I.
PD1+ ABCs
### of CD20+CD21- CD11c+PD-1+ cells
Responder
Non-Responder
J.
CD79a+ ABCs
### of CD79a+CD21- CD11c+cells
K.
CD79a+PD1+ ABCs
### of CD79a+CD21- CD11c+PD-1+cells
L.
PD1+ ABCs
### of CD20+CD21- CD11c+PD-1+ cells
Pre-BCG
Post-BCG
M.
TA-ImmatureTLS
LaminaPropria
Cytotoxic_Tcells
Regulatory_Tcells
Helper_Tcells
CD163+M2-like_cells
CD11c+ DCs
Memory B cells
PD1+ ABCs
ABCs
PD1+ B cells
B cells
100% 50% 0% 50%
Responder
NonResponder_PreBCG
NonResponder_PostBCG
N.
CD79a- ABCs
### of CD79a- CD20-CD11c+ cells
Responder
Non-Responder

***Figure S2. Spatial characterization of myeloid and T cell subsets in the NMIBC tumor microenvironment.*** Bar plots showing the spatial abundance of $CD79a^{-}CD20^{-}CD11c^{+}$ cells (**A**) and $CD79a^{-}CD20^{-}CD11c^{-}CD163^{+}$ cells (**B**) across different compartments including tumor-associated (TA) stroma, mature TA-tertiary lymphoid structures (TLS), immature TA-TLS, and lamina propria (LP) for pre- and post-BCG samples. Representative whole-section multiplex immunofluorescence (mIF) images illustrating spatial organization of T cell subsets across annotated tissue regions, including mature TA-TLS, TA-stroma, immature TA-TLS, and lamina propria (**C**). Bar plots showing quantification of $CD3^{+}$ total T cells (**D**), $CD3^{+}CD4^{+}CD8^{-}$ helper T cells (**E**), $CD3^{+}CD4^{-}CD8^{+}PD\text{-}1^{+}$ T cells (**F**), and $CD3^{+}CD4^{+}CD8^{-}PD\text{-}1^{+}$ T cells (**G**), $CD3^{+}CD4^{+}CD8^{-}$ helper T cells (**H**), $CD3^{+}CD4^{-}CD8^{+}$ cytotoxic T cells (**I**), $CD3^{+}CD4^{+}CD8^{-}PD\text{-}1^{+}$ T cells (**J**), $CD3^{+}CD4^{-}CD8^{+}PD\text{-}1^{+}$ T cells (**K**), and $CD3^{+}CD4^{+}CD8^{-}FoxP3^{+}$ $T_{reg}$ cells (**L**) across tissue compartments in non-responders for different treatment timepoints. Each point represents an average of 3-6 ROIs per sample. Bars indicate mean ± SD. All statistical analyses were performed in GraphPad Prism using non-parametric tests; $*p < 0.05$.

### *Figure S2*

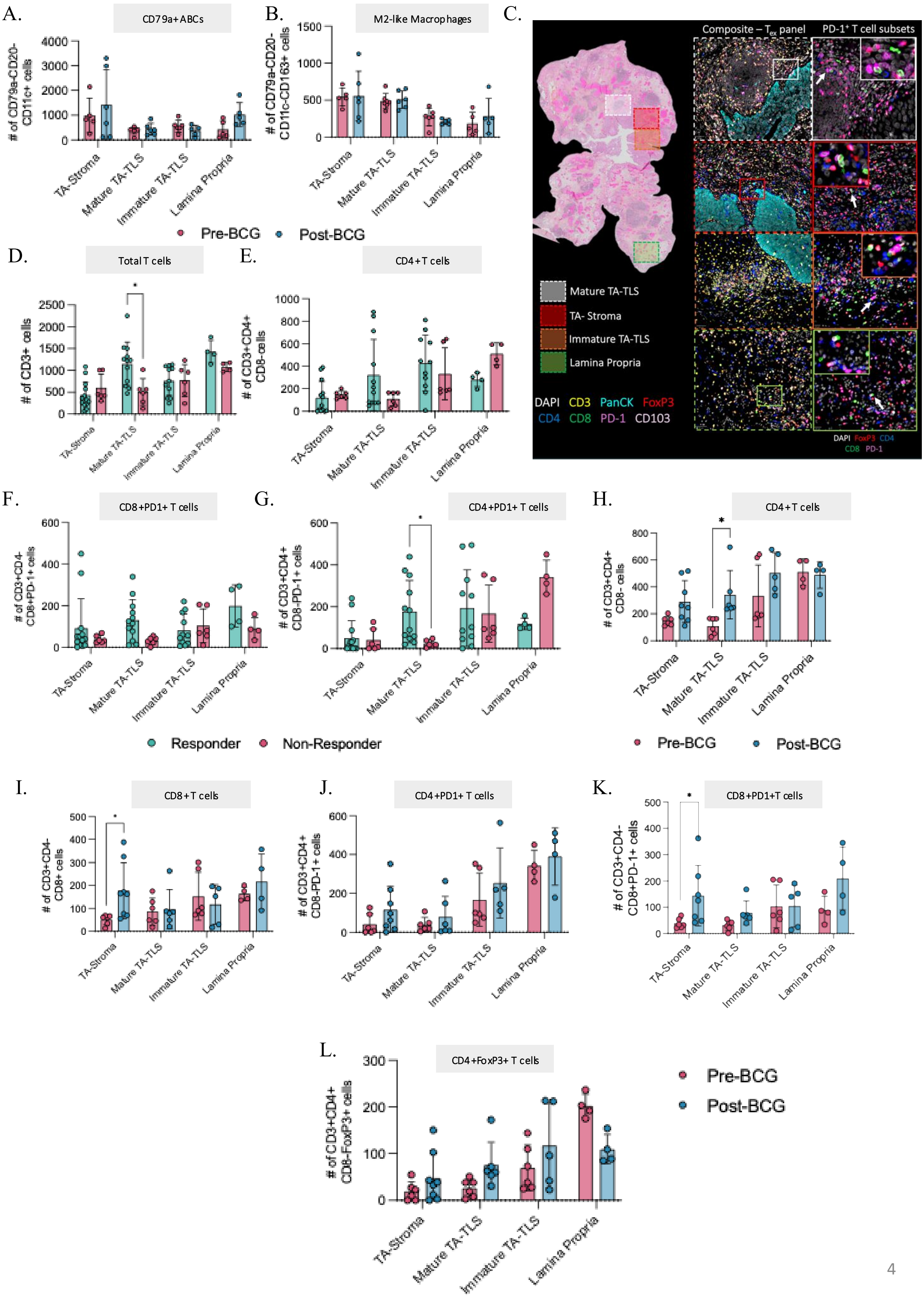
A.
CD79a+ABCs
### of CD79a-CD20-CD11c+ cells
B.
M2-like Macrophages
### of CD79a-CD20-CD11c-CD163+ cells
C.
Composite – Tex panel
PD-1+ T cell subsets
Mature TA-TLS
TA- Stroma
Immature TA-TLS
Lamina Propria
DAPI CD3 PanCK FoxP3
CD4 CD8 PD-1 CD103
Pre-BCG
Post-BCG
D.
Total T cells
### of CD3+ cells
E.
CD4+T cells
### of CD3+CD4+ CD8-cells
F.
CD8+PD1+ T cells
### of CD3+CD4- CD8+PD-1+ cells
G.
CD4+PD1+ T cells
### of CD3+CD4+ CD8-PD-1+ cells
H.
CD4+T cells
### of CD3+CD4+ CD8- cells
Responder
Non-Responder
Pre-BCG
Post-BCG
I.
CD8+T cells
### of CD3+CD4- CD8+ cells
J.
CD4+PD1+ T cells
### of CD3+CD4+ CD8-PD-1+ cells
K.
CD8+PD1+T cells
### of CD3+CD4- CD8+PD-1+ cells
L.
CD4+FoxP3+ T cells
### of CD3+CD4+ CD8-FoxP3+ cells
Pre-BCG
Post-BCG
TA-Stroma
Mature TA-TLS
Immature TA-TLS
Lamina Propria

***Figure S3. Spatial organization and cellular interactions of ABCs in NMIBC tumors associated with BCG response.*** Stacked bar plot showing the distribution of tumor epithelial, B cell, T cell, and myeloid populations, including ABCs, within the bladder TIME across responders, and pre-BCG and post-BCG non-responders (**A**). Ridge plots showing the spatial distribution of ABCs relative to tumor epithelial cells, represented as k-distance to the nearest cancer cell (k = 10), displayed for individual samples and grouped by treatment response with hierarchical clustering illustrating relationships between cell types (ROI S1 and S2) (**B**). Heatmap depicting pairwise spatial distances between major immune populations and tumor epithelium in pre-BCG non-responders **C**). Chord diagram illustrating predicted spatial interactions between ABCs and other immune and tumor cell populations in post-BCG non-responder tumors, highlighting the preferential proximity of ABCs to multiple immune subsets and tumor epithelium (**D**). All analyses were performed in R and visualized using the *ggplot2* package.

*Figure S3*

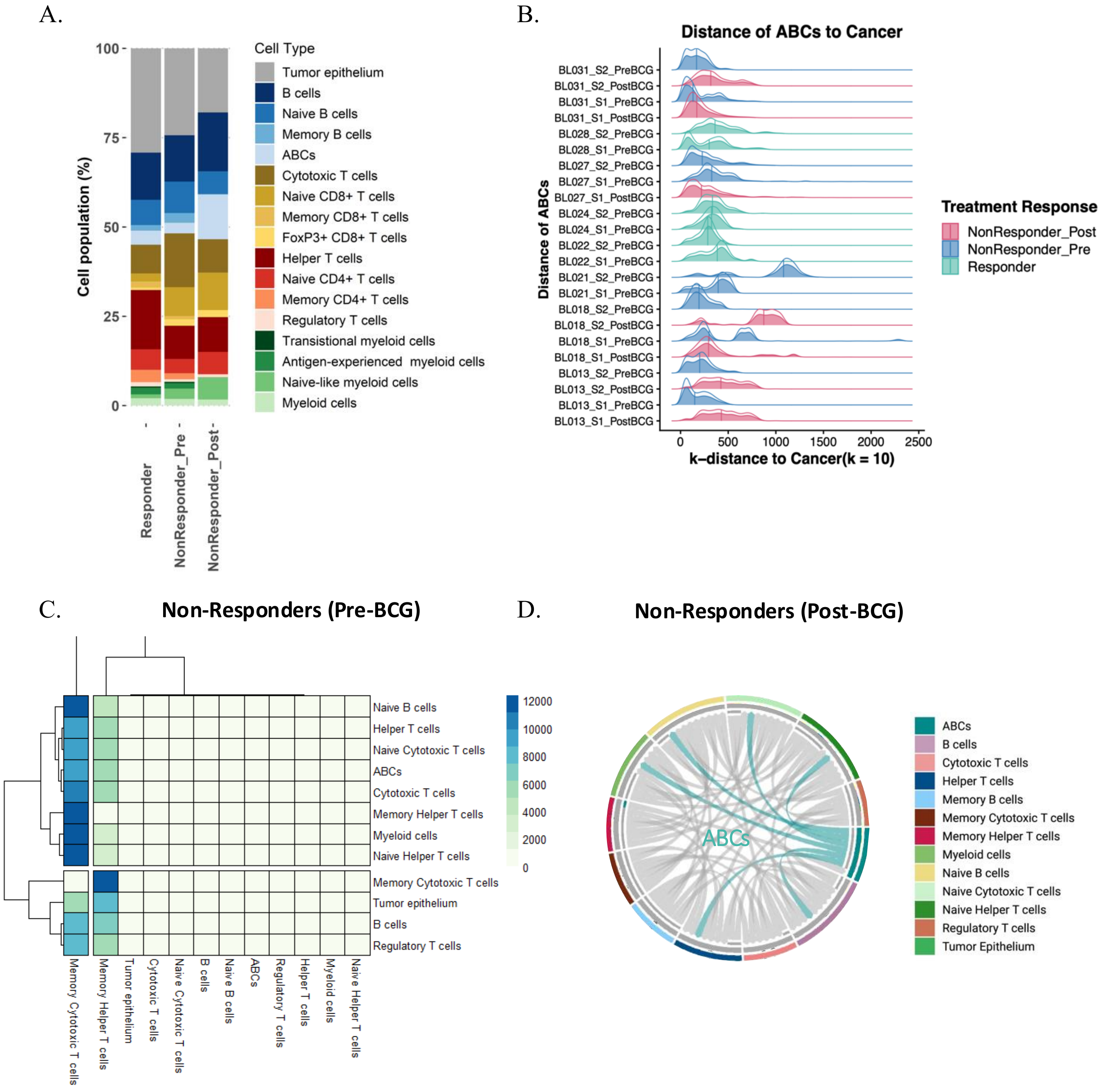
A.
Cell Type
Tumor epithelium
B cells
Naive B cells
Memory B cells
ABCs
Cytotoxic T cells
Naive CD8+ T cells
Memory CD8+ T cells
FoxP3+ CD8+ T cells
Helper T cells
Naive CD4+ T cells
Memory CD4+ T cells
Regulatory T cells
Transitional myeloid cells
Antigen-experienced myeloid cells
Naive-like myeloid cells
Myeloid cells
Cell population (%)
Responder
NonResponder_Pre
NonResponder_Post
B.
Distance of ABCs to Cancer
Distance of ABCs
k–distance to Cancer(k = 10)
Treatment Response
NonResponder_Post
NonResponder_Pre
Responder
C.
Non-Responders (Pre-BCG)
D.
Non-Responders (Post-BCG)
ABCs

***Figure S4. Spatial transcriptomic profiling of NMIBC tumors reveals cellular composition and epithelial state differences associated with BCG response.*** Quality control metrics for Xenium *in situ* spatial transcriptomic data showing distributions of detected features (nFeature_Xenium) and transcript counts (nCount_Xenium) across tissue sections P1-P6 (**A**). UMAP visualization of integrated spatial transcriptomic profiles colored by annotated cell types across pre-BCG responders, and pre-BCG and post-BCG non-responders (**B**). Dot plots illustrating the distribution of cell types within spatially defined cell niches in responders (**C**) and pre-BCG non-responders (**D**); dot size represents the proportion of cells expressing niche-defining markers, and color indicates scaled expression levels. Reactome pathway enrichment analysis comparing responders, to pre-BCG and post-BCG non-responders, highlighting differential activation of immune signaling pathways (**E**). Feature UMAP plots showing spatial expression of canonical B cell markers (*MS4A1, CD79A, CD19*), confirming identification of B cell clusters (**F**). UMAP visualization of B cell and plasma cell subsets identifying ten transcriptionally distinct clusters across pre-BCG responders and both pre- and post-BCG non-responders (**G**). UMAP plots depicting spatial distribution of ten epithelial clusters across treatment groups and timepoints (**H**). Hallmark pathway enrichment analysis of epithelial clusters revealing distinct pathway activation patterns between responders (pre-BCG) and non-responders (pre- and post-BCG) (**I**). All analyses were performed in R and visualized using the *ggplot2* package.

***Figure S4***

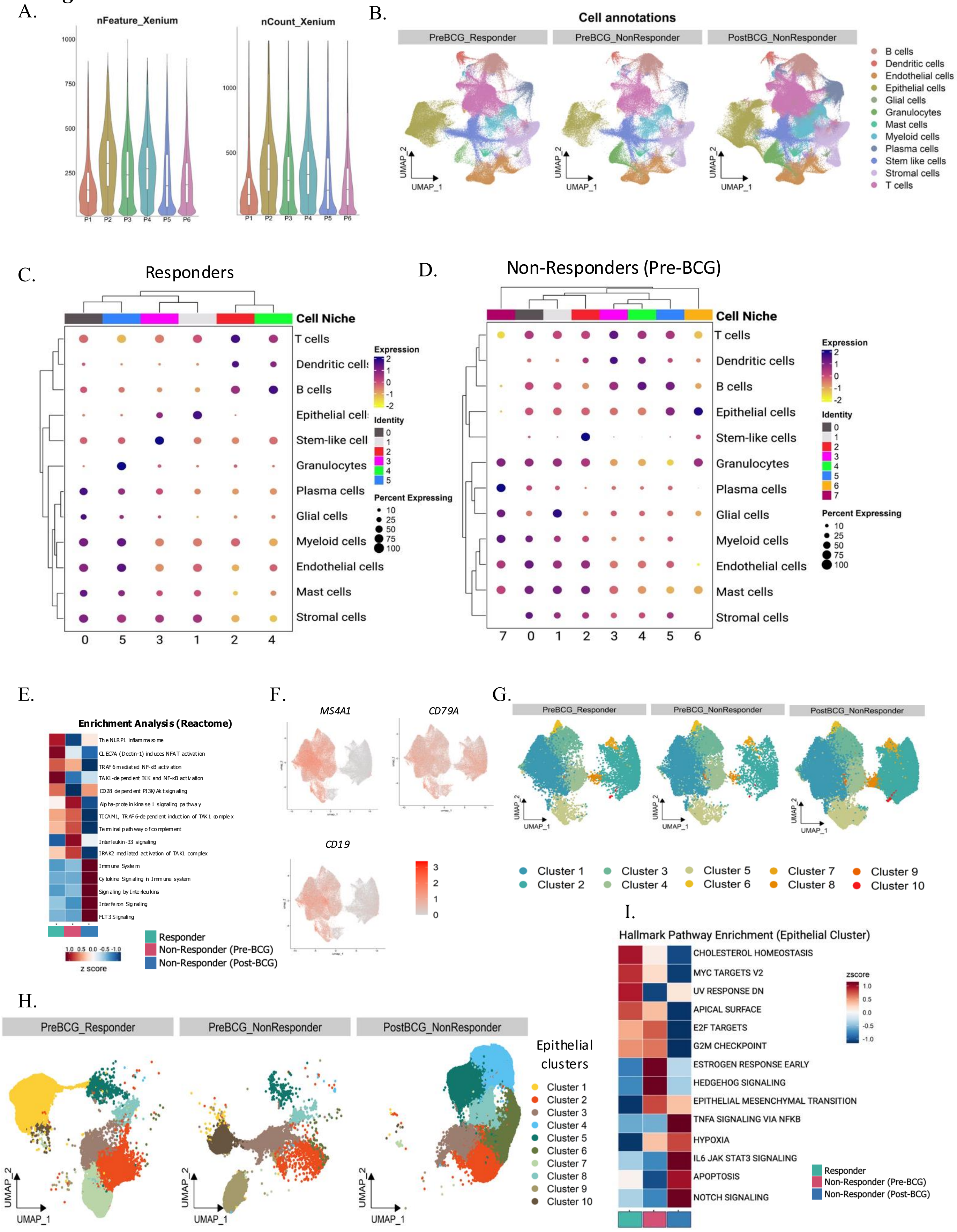

A.
nFeature_Xenium
1000
750
500
250
P1
P2
P3
P4
P5
P6
nCount_Xenium
1000
500
B.
Cell annotations
PreBCG_Responder
PreBCG_NonResponder
PostBCG_NonResponder
UMAP_1
UMAP_2
B cells
Dendritic cells
Endothelial cells
Epithelial cells
Glial cells
Granulocytes
Mast cells
Myeloid cells
Plasma cells
Stem like cells
Stromal cells
T cells
C.
Responders
Cell Niche
T cells
Dendritic cells
B cells
Epithelial cells
Stem-like cell
Granulocytes
Plasma cells
Glial cells
Myeloid cells
Endothelial cells
Mast cells
Stromal cells
0
5
3
1
2
4
Expression
Identity
Percent Expressing
D.
Non-Responders (Pre-BCG)
Cell Niche
T cells
Dendritic cells
B cells
Epithelial cells
Stem-like cells
Granulocytes
Plasma cells
Glial cells
Myeloid cells
Endothelial cells
Mast cells
Stromal cells
7
0
1
2
3
4
5
6
Expression
Identity
Percent Expressing
E.
Enrichment Analysis (Reactome)
The NLRP1 inflammasome
CLEC7A (Dectin-1) induces NFAT activation
TRAF6 mediated NF-κB activation
TAK1-dependent IKK and NF-κB activation
CD28 dependent PI3K/Akt signaling
Alpha-protein kinase 1 signaling pathway
TICAM1, TRAF6-dependent induction of TAK1 complex
Terminal pathway of complement
Interleukin-33 signaling
IRAK2 mediated activation of TAK1 complex
Immune System
Cytokine Signaling in Immune system
Signaling by Interleukins
Interferon Signaling
FLT3 Signaling
z score
Responder
Non-Responder (Pre-BCG)
Non-Responder (Post-BCG)
F.
MS4A1
CD79A
CD19
G.
PreBCG_Responder
PreBCG_NonResponder
PostBCG_NonResponder
UMAP_1
UMAP_2
Cluster 1
Cluster 2
Cluster 3
Cluster 4
Cluster 5
Cluster 6
Cluster 7
Cluster 8
Cluster 9
Cluster 10
H.
PreBCG_Responder
PreBCG_NonResponder
PostBCG_NonResponder
UMAP_1
UMAP_2
Epithelial clusters
Cluster 1
Cluster 2
Cluster 3
Cluster 4
Cluster 5
Cluster 6
Cluster 7
Cluster 8
Cluster 9
Cluster 10
I.
Hallmark Pathway Enrichment (Epithelial Cluster)
CHOLESTEROL HOMEOSTASIS
MYC TARGETS V2
UV RESPONSE DN
APICAL SURFACE
E2F TARGETS
G2M CHECKPOINT
ESTROGEN RESPONSE EARLY
HEDGEHOG SIGNALING
EPITHELIAL MESENCHYMAL TRANSITION
TNFA SIGNALING VIA NFKB
HYPOXIA
IL6 JAK STAT3 SIGNALING
APOPTOSIS
NOTCH SIGNALING
zscore
Responder
Non-Responder (Pre-BCG)
Non-Responder (Post-BCG)

***Figure S5. Gating strategy for identifying B cell subsets from peripheral blood of NMIBC patients.*** Hierarchical gating strategy defining circulating $CD19^+$ B cell subsets, including naïve, memory, transitional and ABCs. ABCs were further subclassified based on C27 and IgD expression (**A**). Single cell suspensions from isolated B cells were stained with antibodies and subjected to multispectral flow cytometric analysis. Single cells was first identified by plotting forward scatter width (FSC-Width) against side scatter area (FSC-A). Events deviating from the linear cell population, called as doublets, were excluded. Live cells were evaluated by gating on viability dye-negative cell. CD45 was used to discriminate the total immune cell population. For identifying ABCs, the following gating strategy was applied: $CD19^+$ cells revealed the proportion of total B cells. Further plotting against CD21 and CD11c markers was used to identify ABC population (identified as $CD21^{-/low}$ $CD11c^+$). ABCs were further gated against CD27, IgD, and IgM to delineate ABC subsets. For identification of memory B subsets, the following markers was used: $CD21^{-/+}$ $CD27^+$ $IgM^-$ $IgD^-$ class switched memory B cells, and $CD21^{-/+}$ $CD27^+$ $IgM^+$ $IgD^+$ class un-switched memory B cells **(B).** Single color (SC) and fluorescence minus one (FMO) was used to determine the position of gating for all subsets and further compared with unstained samples. All analysis were done in FlowJo® software.

*Figure S5*

A.

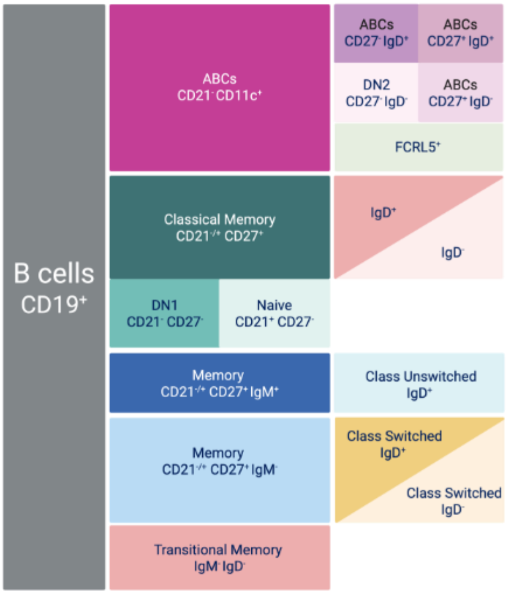
B cells
CD19+
ABCs
CD21- CD11c+
ABCs
CD27- IgD+
ABCs
CD27+ IgD+
DN2
CD27- IgD-
ABCs
CD27+ IgD-
FCRL5+
Classical Memory
CD21-/+ CD27+
IgD+
IgD-
DN1
CD21- CD27-
Naive
CD21+ CD27-
Memory
CD21-/+ CD27+ IgM+
Class Unswitched
IgD+
Memory
CD21-/+ CD27+ IgM-
Class Switched
IgD+
Class Switched
IgD-
Transitional Memory
IgM- IgD-

B.

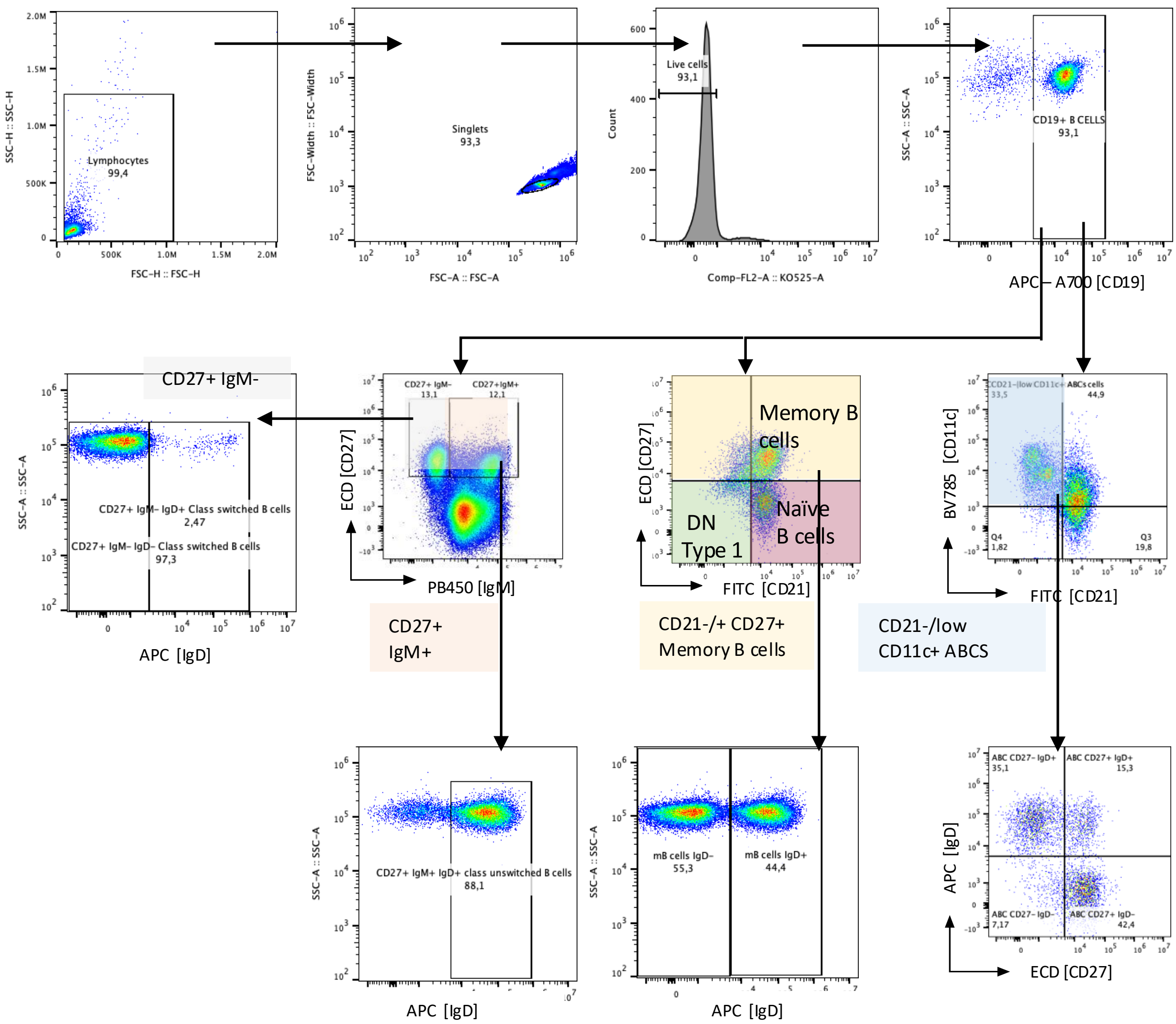
SSC-H :: SSC-H
FSC-H :: FSC-H
Lymphocytes
99,4
FSC-Width :: FSC-Width
FSC-A :: FSC-A
Singlets
93,3
Count
Comp-FL2-A :: KO525-A
Live cells
93,1
SSC-A :: SSC-A
APC – A700 [CD19]
CD19+ B CELLS
93,1
CD27+ IgM-
CD27+ IgM- IgD+ Class switched B cells
2,47
CD27+ IgM- IgD- Class switched B cells
97,3
APC [IgD]
ECD [CD27]
PB450 [IgM]
CD27+ IgM-
13,1
CD27+IgM+
12,1
CD27+
IgM+
Memory B
cells
DN
Type 1
Naïve
B cells
FITC [CD21]
CD21-/+ CD27+
Memory B cells
BV785 [CD11c]
CD21-/low CD11c+
33,5
ABCs cells
44,9
Q4
1,82
Q3
19,8
CD21-/low
CD11c+ ABCS
CD27+ IgM+ IgD+ class unswitched B cells
88,1
mB cells IgD-
55,3
mB cells IgD+
44,4
APC [IgD]
ABC CD27- IgD+
35,1
ABC CD27+ IgD+
15,3
ABC CD27- IgD-
7,17
ABC CD27+ IgD-
42,4

***Figure S6*. *Profiling of circulating B cell subsets in response to BCG across different timepoints*.** Bar graph illustrates the viability (**A**) and purity of $CD19^+$ B cells (**B**) after isolation from NMIBC patients undergoing BCG immunotherapy. Spearman correlation analysis plot demonstrating lack of significant correlation of ABCs frequencies (*r = 0.08865, p = ns*) between pre-BCG and post-$4^{th}$ BCG timepoints in BCG responders (**C**). Bar graph showing the frequency change of ABCs ($CD19^+CD21^-CD11c^+$; **D**), DN1 B cells ($CD19^+CD21^-CD27^-$; **(E & F**), DN2 B cells ($CD19^+CD27^-IgD^-$; **(G & H)** $CD27^+IgD^-$ ABCs (**I** & **J**), $CD27^+IgD^+$ ABCs (**K**), $CD27^-IgD^+$ ABCs (**L**), and $CD27^-IgD^-$ $FCRL5^+$ ABCs (**M**) in responders (R) and non-responders (NR) across different timepoints (pre-BCG, post-$1^{st}$ BCG, and post-$4^{th}$ BCG). Representative t-SNE plots of 4 down samples (10,000 $CD19^+$ B cells) and concatenated samples showing flow cytometry-based clustering of B cell populations in responders and non-responders before and after BCG therapy (**N**). Bar graphs showing the frequency change of class-switched B cells (**O & P**), class-switched Memory B cells in responders (**Q**), and naïve B cells in responders (**R**). Bar graphs showing the frequency change of class-switched memory B cells in non-responders (**S**), unswitched memory B cells in non-responders **(T)** vs responders (**U**). Analysis of flow cytometry data was performed using FlowJo® software. All statistical analysis was performed using GraphPad Prism. Non-parametric Kruskal-Wallis test with Dunn's multiple comparison tests was applied to determine statistical significance between groups (*p<0.05). Bar graphs represent mean ± SD. Frequencies in bar graph are shown as percentage change per sample across timepoints. R: Responder (n = 22) | NR: Non-Responder (n = 9).

*Figure S6*

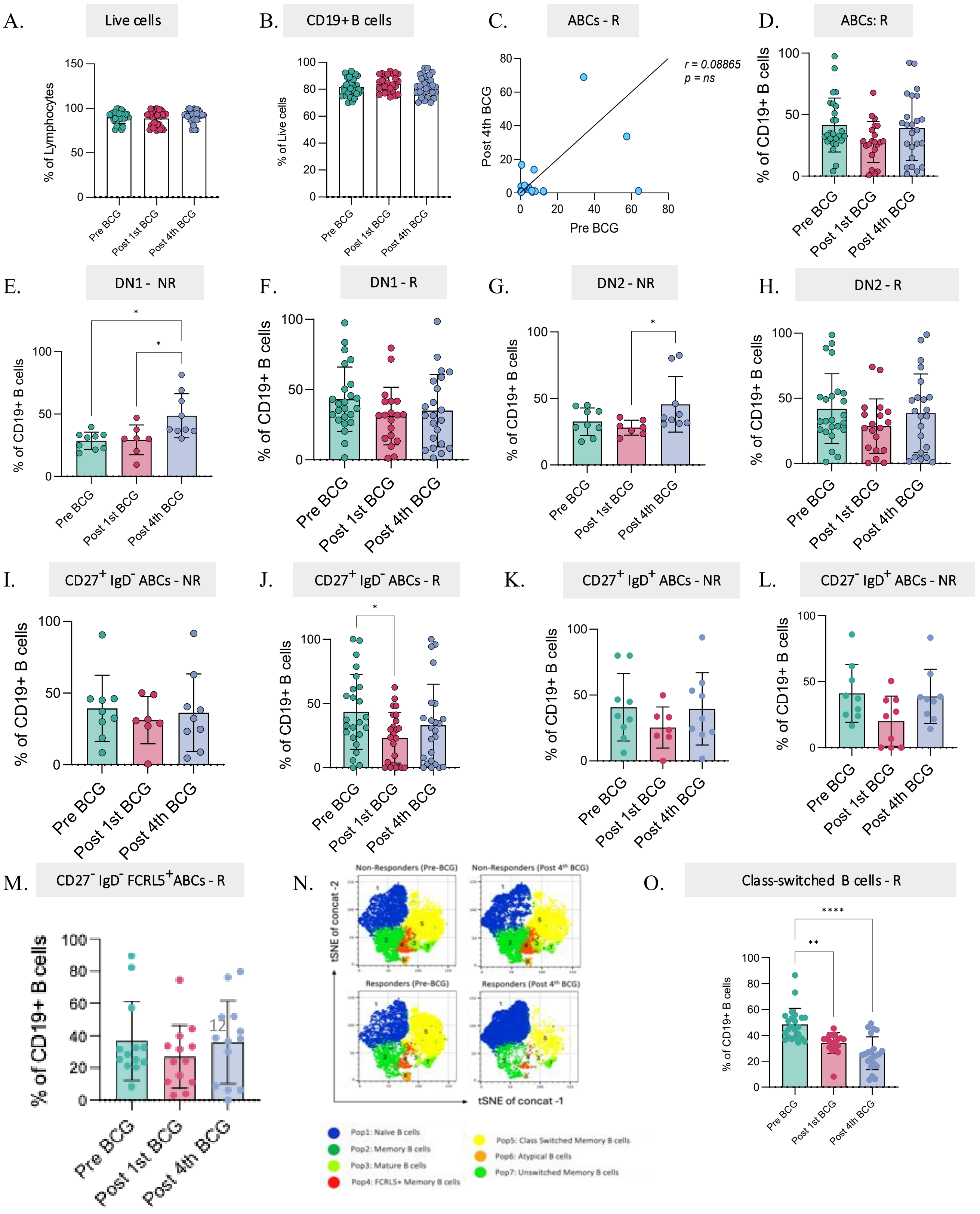

A.
Live cells
% of Lymphocytes
Pre BCG
Post 1st BCG
Post 4th BCG
B.
CD19+ B cells
% of Live cells
C.
ABCs - R
r = 0.08865
p = ns
Post 4th BCG
Pre BCG
D.
ABCs: R
% of CD19+ B cells
E.
DN1 - NR
F.
DN1 - R
G.
DN2 - NR
H.
DN2 - R
I.
CD27+ IgD- ABCs - NR
J.
CD27+ IgD- ABCs - R
K.
CD27+ IgD+ ABCs - NR
L.
CD27- IgD+ ABCs - NR
M.
CD27- IgD- FCRL5+ABCs - R
N.
Non-Responders (Pre-BCG)
Non-Responders (Post 4th BCG)
Responders (Pre-BCG)
Responders (Post 4th BCG)
tSNE of concat -2
tSNE of concat -1
Pop1: Naïve B cells
Pop2: Memory B cells
Pop3: Mature B cells
Pop4: FCRL5+ Memory B cells
Pop5: Class Switched Memory B cells
Pop6: Atypical B cells
Pop7: Unswitched Memory B cells
O.
Class-switched B cells - R
% of CD19+ B cells

*Figure S6 continued…*

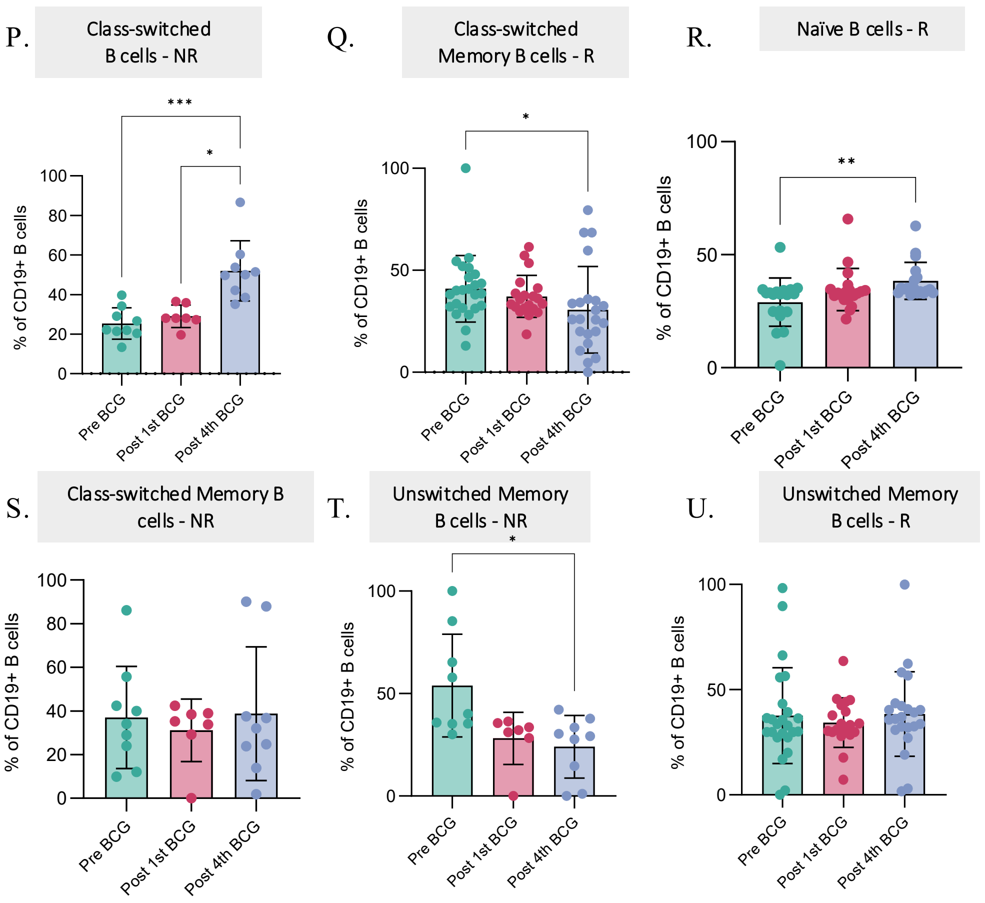

***Figure S7. Single-cell transcriptomic profiling and pseudotime dynamics of B cell subsets.*** Bar graph depicting the mean RNA counts and Features per sample in the scRNAseq data (**A**). Violin plot illustrating the nFeature, nCount and mitochondrial content in individual samples after pre-processing of scRNA seq data (**B**). Feature plot representing expression of pan-B cell markers (*CD19, MS4A1, CD79A, CD79B)* in the B cell identified clusters shown as UMAP (**C**). Violin plot depicting distribution and expression of canonical B cell markers (*TCL1A, CD27,CD38, XBP1*) and other cluster defining genes *(FCER1A, FCRL5, ITGAX, TBX21, CCR7, SLC38A11, EFHD2, GNLY)* in different B cell clusters (**D**). UMAP depicting the distribution of different B cell clusters in each sample **(E)**. Dot plot illustrating statistically significant ligand receptor interaction pairs enriched in ABCs. Dot size reflects significance of communication probability; colour intensity (red to blue) reflects interaction strength (**F**). Analysis of scRNAseq data was performed using Seurat package in R. All statistical analysis was performed using R. S001A, S002A, S003A, S004A represent the pre-BCG samples | S001B, S002B, S003B, S004B represent the post-4$^{th}$ BCG samples.

*Figure S7*

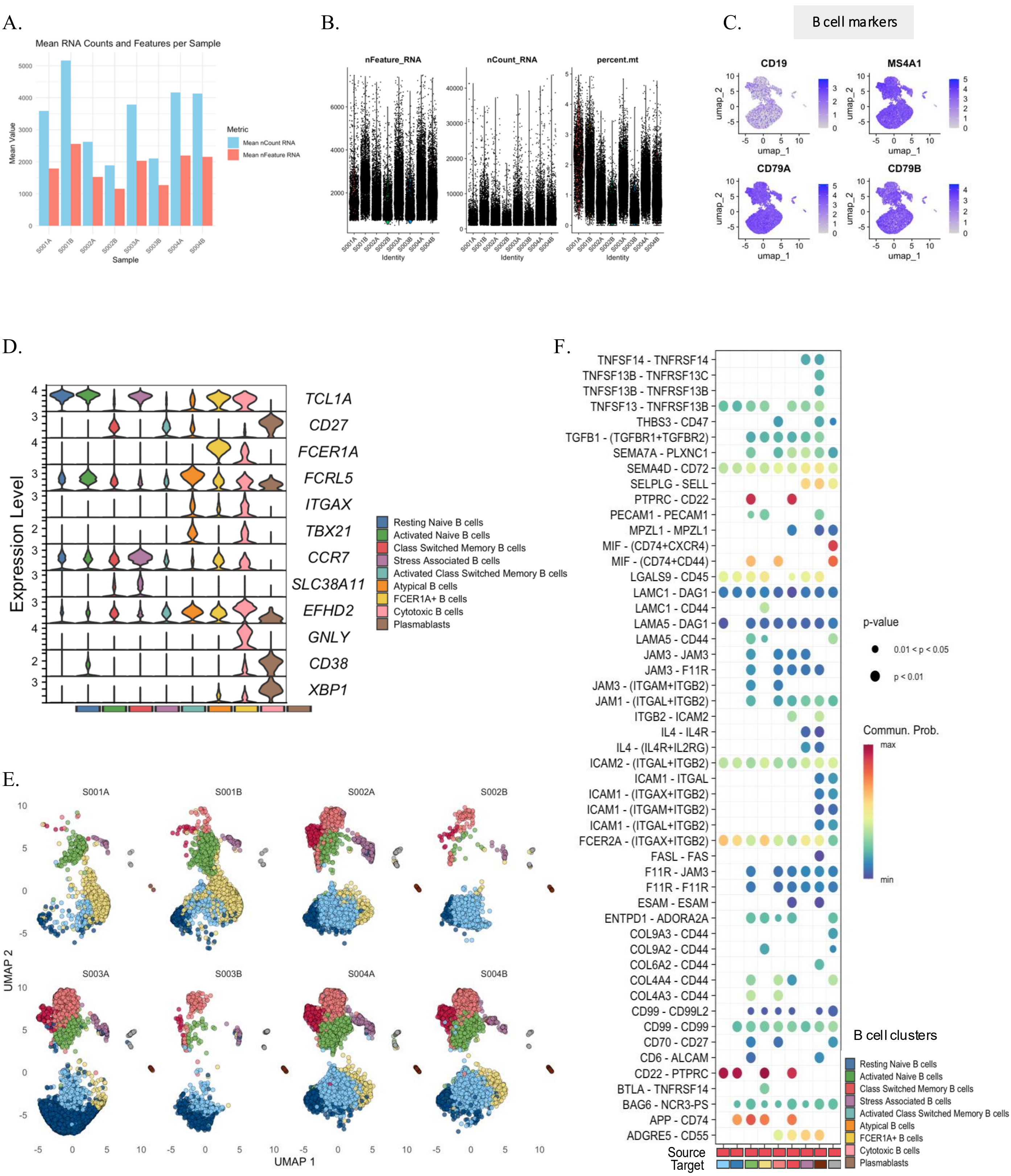
A.
Mean RNA Counts and Features per Sample
Mean Value
Sample
Metric
Mean nCount RNA
Mean nFeature RNA
S001A
S001B
S002A
S002B
S003A
S003B
S004A
S004B
B.
nFeature_RNA
nCount_RNA
percent.mt
Identity
C.
B cell markers
CD19
MS4A1
CD79A
CD79B
umap_1
umap_2
D.
Expression Level
TCL1A
CD27
FCER1A
FCRL5
ITGAX
TBX21
CCR7
SLC38A11
EFHD2
GNLY
CD38
XBP1
Resting Naive B cells
Activated Naive B cells
Class Switched Memory B cells
Stress Associated B cells
Activated Class Switched Memory B cells
Atypical B cells
FCER1A+ B cells
Cytotoxic B cells
Plasmablasts
E.
UMAP 1
UMAP 2
F.
TNFSF14 - TNFRSF14
TNFSF13B - TNFRSF13C
TNFSF13B - TNFRSF13B
TNFSF13 - TNFRSF13B
THBS3 - CD47
TGFB1 - (TGFBR1+TGFBR2)
SEMA7A - PLXNC1
SEMA4D - CD72
SELPLG - SELL
PTPRC - CD22
PECAM1 - PECAM1
MPZL1 - MPZL1
MIF - (CD74+CXCR4)
MIF - (CD74+CD44)
LGALS9 - CD45
LAMC1 - DAG1
LAMC1 - CD44
LAMA5 - DAG1
LAMA5 - CD44
JAM3 - JAM3
JAM3 - F11R
JAM3 - (ITGAM+ITGB2)
JAM1 - (ITGAL+ITGB2)
ITGB2 - ICAM2
IL4 - IL4R
IL4 - (IL4R+IL2RG)
ICAM2 - (ITGAL+ITGB2)
ICAM1 - ITGAL
ICAM1 - (ITGAX+ITGB2)
ICAM1 - (ITGAM+ITGB2)
ICAM1 - (ITGAL+ITGB2)
FCER2A - (ITGAX+ITGB2)
FASL - FAS
F11R - JAM3
F11R - F11R
ESAM - ESAM
ENTPD1 - ADORA2A
COL9A3 - CD44
COL9A2 - CD44
COL6A2 - CD44
COL4A4 - CD44
COL4A3 - CD44
CD99 - CD99L2
CD99 - CD99
CD70 - CD27
CD6 - ALCAM
CD22 - PTPRC
BTLA - TNFRSF14
BAG6 - NCR3-PS
APP - CD74
ADGRE5 - CD55
Source
Target
p-value
0.01 < p < 0.05
p < 0.01
Commun. Prob.
max
min
B cell clusters
Resting Naive B cells
Activated Naive B cells
Class Switched Memory B cells
Stress Associated B cells
Activated Class Switched Memory B cells
Atypical B cells
FCER1A+ B cells
Cytotoxic B cells
Plasmablasts

***Figure S8*****. Cell-cell communication networks and signaling pattern analysis across B cell subsets.** Global interaction network depicting the overall strength of inferred ligand-receptor interactions among annotated B cell clusters. Edge thickness reflects interaction strength, and node size corresponds to the total communication probability for each cluster (**A**). Chord diagrams summarizing all inferred ligand-receptor interactions across B cell cluster, focusing on all the receptors (**B**) and ligands (**C**) in the ABC cluster. Network representation of FCER2A-mediated signaling pathways, including FCER2A-ITGAX (ITGB2/CD99; **D**) and FCER2A-CR2 (CD99; **E**), highlighting cluster-specific communication patterns. Line plots comparing incoming (**F**) and outgoing (**G**) signaling pattern robustness across increasing numbers of inferred communication patterns, evaluated using Cophenetic and Silhouette measures. Sankey diagram illustrating incoming communication patterns of target B cell populations, linking cell groups to dominant signaling patterns and associated ligand-receptor pathways (**H**). Sankey diagram depicting outgoing communication patterns of secreting B cell populations, connecting source cell groups to signaling and downstream pathways (**I**). All scRNAseq analyses and statistical analyses were performed using the Seurat package in R. Cell-cell communication network analysis was done using CellChatDB in R.

*Figure S8*

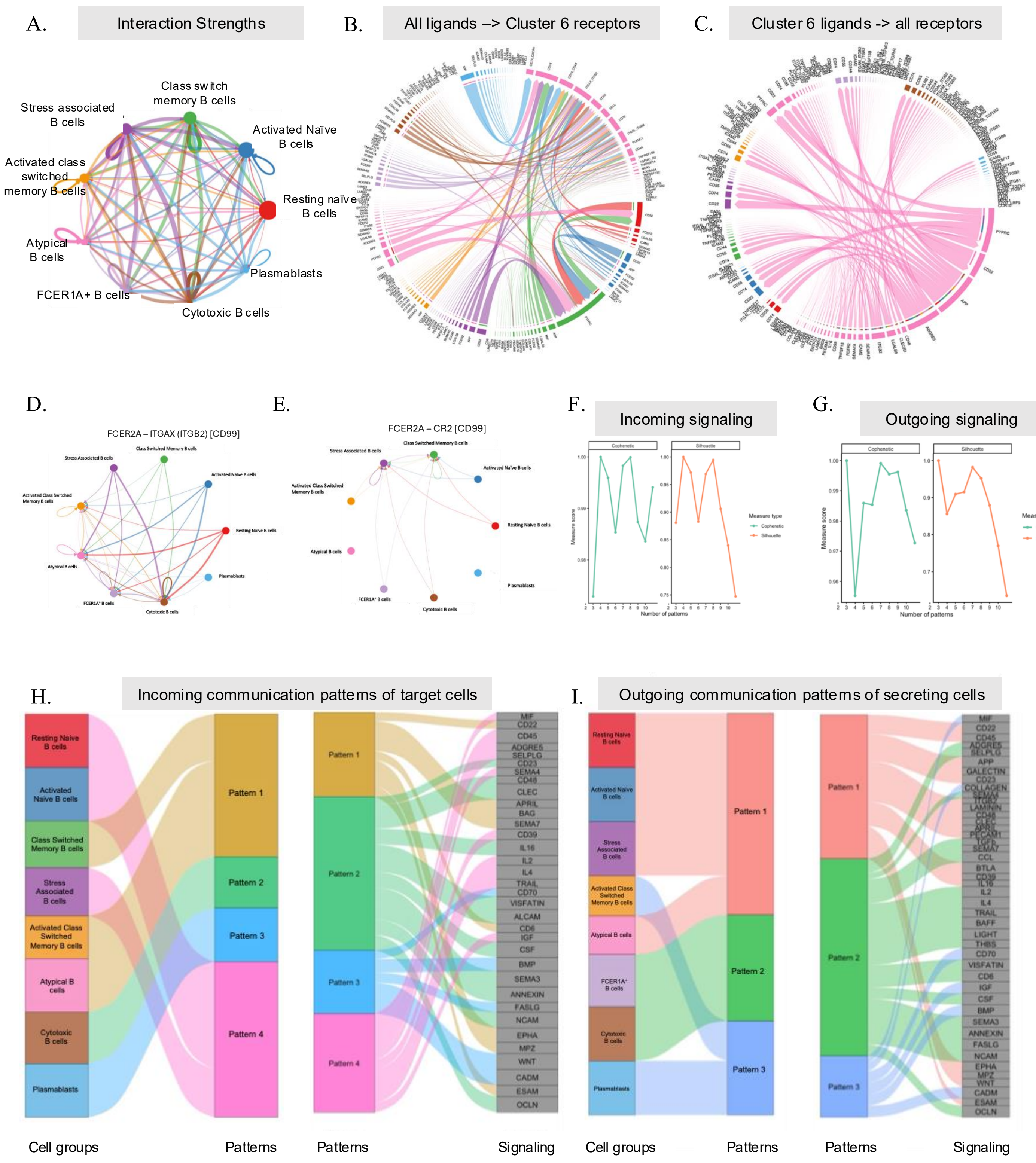
A.
Interaction Strengths
Class switch memory B cells
Stress associated B cells
Activated Naïve B cells
Activated class switched memory B cells
Resting naïve B cells
Atypical B cells
Plasmablasts
FCER1A+ B cells
Cytotoxic B cells
B.
All ligands --> Cluster 6 receptors
C.
Cluster 6 ligands -> all receptors
D.
FCER2A – ITGAX (ITGB2) [CD99]
E.
FCER2A – CR2 [CD99]
F.
Incoming signaling
G.
Outgoing signaling
Cophenetic
Silhouette
Measure type
Measure score
Number of patterns
H.
Incoming communication patterns of target cells
Resting Naive B cells
Activated Naive B cells
Class Switched Memory B cells
Stress Associated B cells
Activated Class Switched Memory B cells
Atypical B cells
Cytotoxic B cells
Plasmablasts
Pattern 1
Pattern 2
Pattern 3
Pattern 4
I.
Outgoing communication patterns of secreting cells
Cell groups
Patterns
Patterns
Signaling
Cell groups
Patterns
Patterns
Signaling

***Figure S9*****. *****Humoral immune profiling and autoantibody landscape associated with BCG response in NMIBC.*** Scatter bar plots showing circulating concentrations of IgG1 (**A**), IgG3 (**B**), IgG4 (**C**), IgA (**D**), and IgM (**E**) in plasma, measured prior to BCG initiation (pre-BCG) and post-4$^{th}$ BCG (post-BCG) in responders and non-responders. Each point represents an individual patient sample. Principal component analysis (PCA) of plasma immunoglobulin profiles demonstrating overall variation across samples stratified by treatment condition and treatment response (**F**). Summary bar plot showing the number of significantly altered IgG-reactive autoantigens detected across the indicated comparisons, highlighting differential humoral reactivity associated with treatment exposure and response status (**G**). Dot plot depicting KEGG (**H**) and GO Biological Process (**I**) pathway enrichment for IgG-reactive antigens in non-responders and responders, respectively, following BCG treatment (paired post-BCG vs pre-BCG comparison). Scatter line plots summarizing ELISA-based quantification of plasma autoreactivity against BCG antigen across serial dilutions (1:100 – 1:100,000) measured longitudinally in non-responders (**J**) and responders (**K**) at multiple time points. Scatter bar plots depict BCG-specific IgG reactivity in the plasma samples, with each dot depicting individual responder samples at 1:100 (**L**) and 1:1,000 dilution (**M**) and non-responder samples at 1:1,000 dilution (**N**). Lines connect longitudinal samples from the same patient. Stacked bar plots showing the distribution of IgA and IgG expression scores within tumor-associated (TA) stroma stratified by response status and treatment timepoint (**O**). Quality control metrics for Olink-based immuno-oncology proteomic profiling, including a scatter plot of sample median signal versus overall assay intensity demonstrating consistent assay performance across samples (**P**) and boxplots of normalized protein expression (NPX) values indicating uniform signal distributions and absence of major technical outliers (**Q**). All statistical analyses were performed in GraphPad Prism using non-parametric paired t tests and Kruskal–Wallis tests with Dunn's multiple comparison tests; *p<0.05.

*Figure S9*

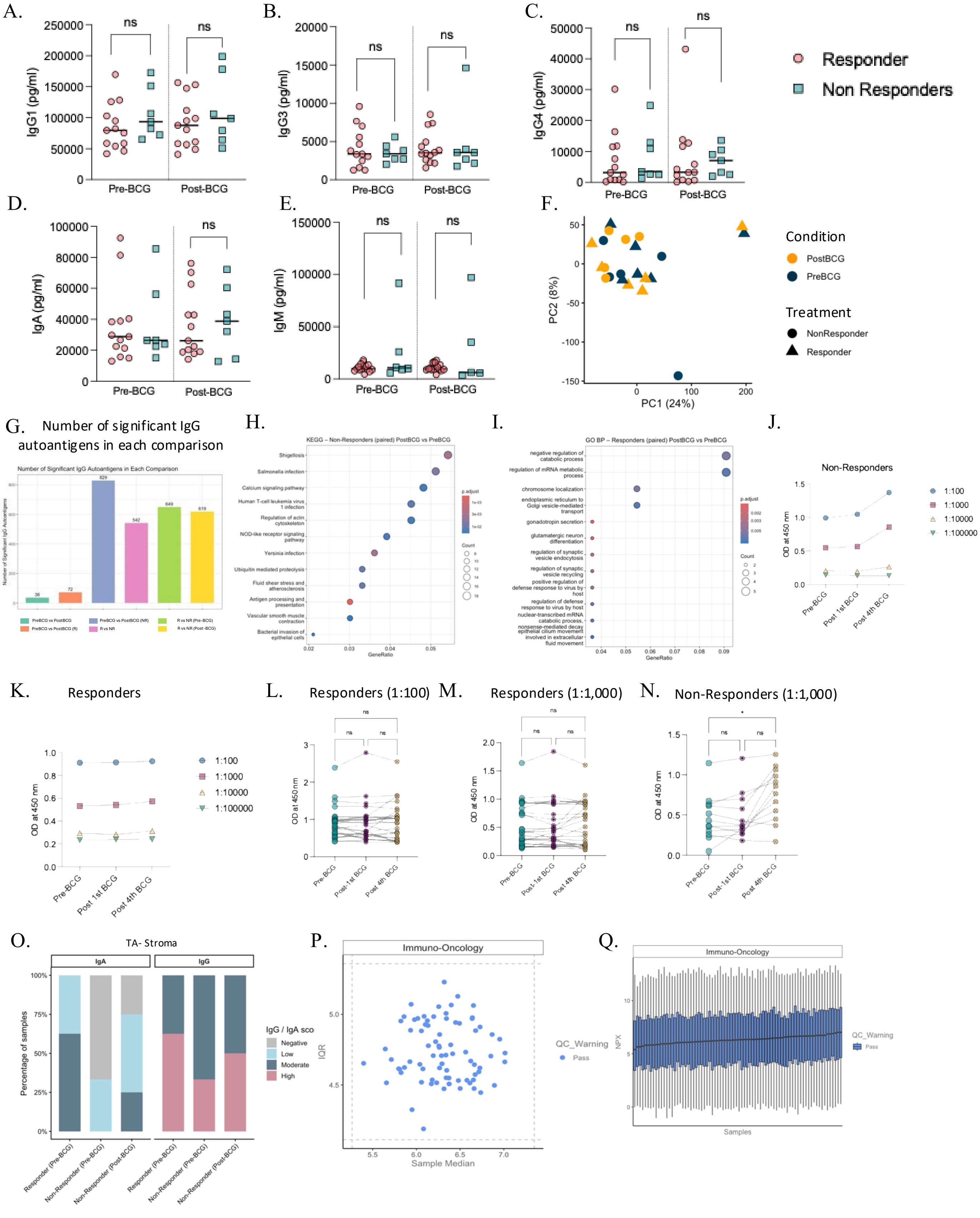

A.
IgG1 (pg/ml)
ns
Pre-BCG
Post-BCG
B.
IgG3 (pg/ml)
C.
IgG4 (pg/ml)
Responder
Non Responders
D.
IgA (pg/ml)
E.
IgM (pg/ml)
F.
PC1 (24%)
PC2 (8%)
Condition
PostBCG
PreBCG
Treatment
NonResponder
Responder
G. Number of significant IgG autoantigens in each comparison
Number of Significant IgG Autoantigens in Each Comparison
H.
KEGG – Non-Responders (paired) PostBCG vs PreBCG
Shigellosis
Salmonella infection
Calcium signaling pathway
Human T-cell leukemia virus 1 infection
Regulation of actin cytoskeleton
NOD-like receptor signaling pathway
Yersinia infection
Ubiquitin mediated proteolysis
Fluid shear stress and atherosclerosis
Antigen processing and presentation
Vascular smooth muscle contraction
Bacterial invasion of epithelial cells
GeneRatio
I.
GO BP – Responders (paired) PostBCG vs PreBCG
negative regulation of catabolic process
regulation of mRNA metabolic process
chromosome localization
endoplasmic reticulum to Golgi vesicle-mediated transport
gonadotropin secretion
glutamatergic neuron differentiation
regulation of synaptic vesicle endocytosis
regulation of synaptic vesicle recycling
positive regulation of defense response to virus by host
regulation of defense response to virus by host
nuclear-transcribed mRNA catabolic process, nonsense-mediated decay
epithelial cilium movement involved in extracellular fluid movement
GeneRatio
J.
Non-Responders
OD at 450 nm
1:100
1:1000
1:10000
1:100000
Pre-BCG
Post 1st BCG
Post 4th BCG
K.
Responders
L.
Responders (1:100)
M.
Responders (1:1,000)
N.
Non-Responders (1:1,000)
O.
TA- Stroma
IgA
IgG
Percentage of samples
IgG / IgA sco
Negative
Low
Moderate
High
Responder (Pre-BCG)
Non-Responder (Pre-BCG)
Non-Responder (Post-BCG)
P.
Immuno-Oncology
IQR
Sample Median
QC_Warning
Pass
Q.
Immuno-Oncology
NPX
Samples
QC_Warning
Pass

***Figure S10. Longitudinal plasma proteomic profiling reveals immune signaling changes associated with BCG treatment in NMIBC.*** Volcano plot depicting differentially abundant plasma proteins following BCG therapy (pre-BCG vs post-4$^{th}$ BCG) across the cohort (**A**). Volcano plot illustrating differential plasma protein abundance at baseline (pre-BCG) between responders and non-responders (**B**). Box plots summarizing longitudinal changes in selected immune mediators in responders (**C**) and non-responders (**D**) comparing pre-BCG and post-4$^{th}$ BCG timepoints. Normalized protein expression (NPX) values are shown for representative proteins (CCL23, CD5, GZMA, TNFRSF21 and IL10) identified in the differential analysis. Dot plot showing KEGG pathway enrichment **(E)** and Gene Ontology (GO) Biological Process enrichment (**F**) for significantly altered proteins between pre-BCG and post-4$^{th}$ BCG samples. Heatmaps showing pairwise correlation matrices of plasma immune-oncology proteins measured at pre-BCG and post-4$^{th}$ BCG in non-responders (**G**) and responders (**H**). Color scale represents Pearson correlation coefficients ranging from negative (blue) to positive (yellow) correlations. Red boxes highlight selected cytokine demonstrating altered correlation structure following BCG therapy. All proteomic measurements were performed using the Olink platform. Data normalization and comparative statistical analyses were conducted in R using non-parametric tests with multiple testing correction where appropriate. *$p<0.05$, **$p<0.01$.

*Figure S10*

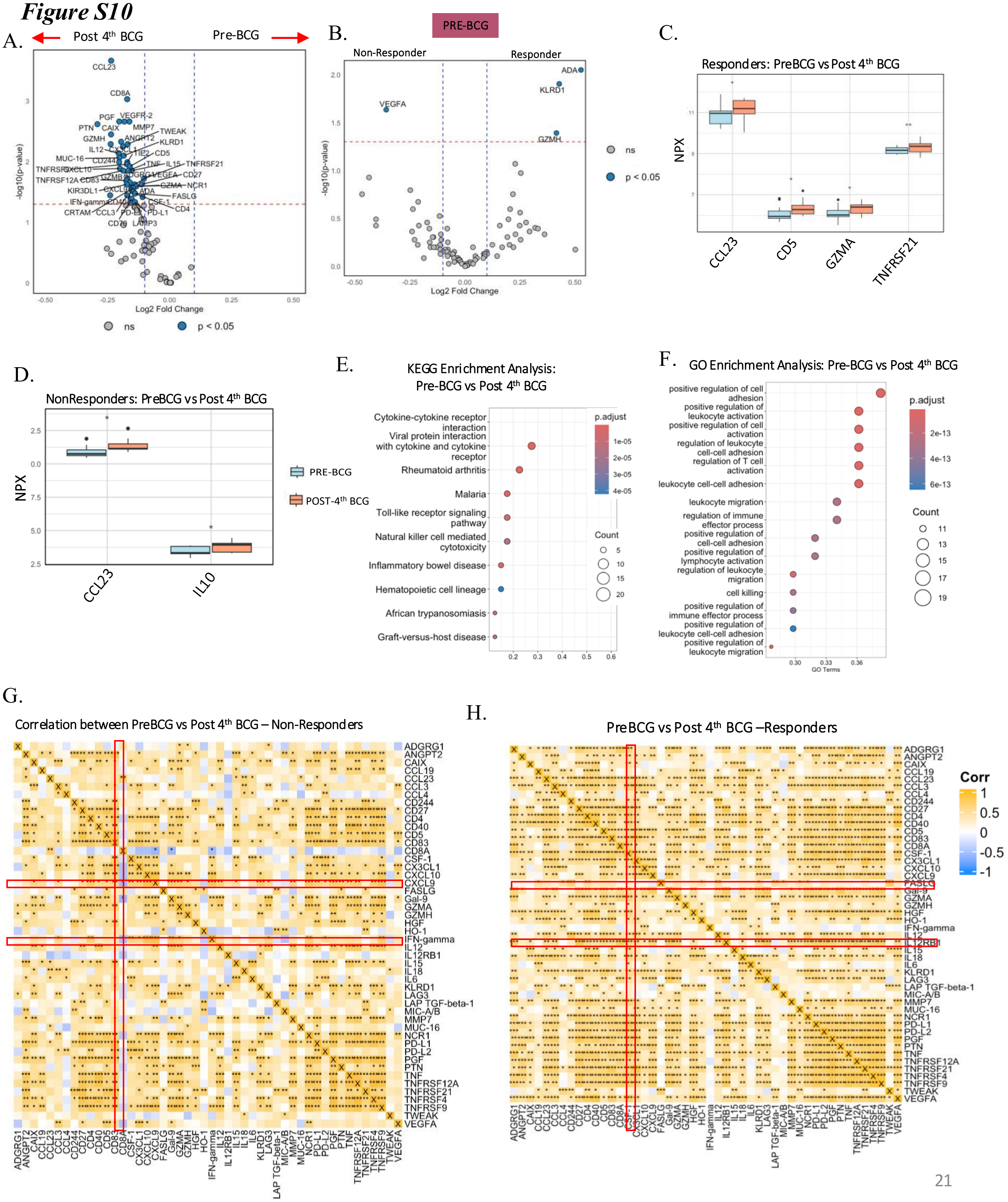

A.
Post 4th BCG
Pre-BCG
-log10(p-value)
Log2 Fold Change
ns
p < 0.05
B.
PRE-BCG
Non-Responder
Responder
ADA
KLRD1
VEGFA
GZMH
-log10(p-value)
Log2 Fold Change
ns
p < 0.05
C.
Responders: PreBCG vs Post 4th BCG
NPX
CCL23
CD5
GZMA
TNFRSF21
D.
NonResponders: PreBCG vs Post 4th BCG
NPX
CCL23
IL10
PRE-BCG
POST-4th BCG
E.
KEGG Enrichment Analysis:
Pre-BCG vs Post 4th BCG
Cytokine-cytokine receptor interaction
Viral protein interaction with cytokine and cytokine receptor
Rheumatoid arthritis
Malaria
Toll-like receptor signaling pathway
Natural killer cell mediated cytotoxicity
Inflammatory bowel disease
Hematopoietic cell lineage
African trypanosomiasis
Graft-versus-host disease
p.adjust
Count
F.
GO Enrichment Analysis: Pre-BCG vs Post 4th BCG
positive regulation of cell adhesion
positive regulation of leukocyte activation
positive regulation of cell activation
regulation of leukocyte cell-cell adhesion
regulation of T cell activation
leukocyte cell-cell adhesion
leukocyte migration
regulation of immune effector process
positive regulation of cell-cell adhesion
positive regulation of lymphocyte activation
regulation of leukocyte migration
cell killing
positive regulation of immune effector process
positive regulation of leukocyte cell-cell adhesion
positive regulation of leukocyte migration
GO Terms
p.adjust
Count
G.
Correlation between PreBCG vs Post 4th BCG – Non-Responders
H.
PreBCG vs Post 4th BCG –Responders
Corr
